# Photo-click Decellularized Matrix Hydrogels for Generating Pancreatic Ductal Organoids

**DOI:** 10.64898/2026.02.16.706185

**Authors:** Ngoc Ha Luong, Kunming Shao, Van Thuy Duong, Xiaoping Bao, Chien-Chi Lin

## Abstract

Pancreatic ductal organoids (PDOs) generated from human induced pluripotent stem cells (iPSCs) can be used to model pancreatic diseases and to conduct drug screening/testing. However, current protocols for generating PDOs rely heavily on tumor-derived Matrigel, which has been shown to upregulate oncogenes. Furthermore, Matrigel has undefined composition and weak mechanical properties that hamper mechanistic studies of cell-material interactions. In this study, we explore photo-clickable decellularized small intestine submucosa-norbornene (dSIS-NB) hydrogels as a Matrigel replacement for generating human iPSC-derived PDOs. To achieve this, pancreatic progenitors (PP) were first differentiated in conventional two-dimensional (2D) culture, aggregated into spheroids, then encapsulated and differentiated within dSIS-NB hydrogels with tunable stiffness. The differentiated organoids were analyzed by morphology, expression of key pancreatic ductal markers, and single-cell RNA sequencing (scRNA-seq). Post-differentiation, PDOs generated in stiffer photo-clickable dSIS-NB hydrogels (shear moduli ∼2.5 kPa) maintained ductal epithelial phenotype and exhibited pronounced forskolin-induced swelling. In contrast, differentiation of PP spheroids in softer dSIS-NB gels (shear moduli ∼0.9 kPa) and Matrigel resulted in a persistent mesenchymal phenotype and failed to generate functional PDOs. Finally, scRNA-seq results revealed that stiffer dSIS-NB hydrogels strongly biased ductal cell differentiation, yielding greater than 97% ductal progeny.

## 1. Introduction

The pancreatic duct transports digestive enzymes,^1^ and its dysfunction contributes to chronic pancreatitis,^2^ cystic fibrosis,^3^ and pancreatic ductal adenocarcinoma (PDAC).^4^ Studying pancreatic ductal diseases in animal models may not fully recapitulate human anatomy and physiology. The gap between animal models and human physiology can be bridged by New Approach Methodologies (NAM), which are more ethical, scalable, and human-relevant.^5^ One promising NAM is stem cell-derived organoids, which have increasingly been used in developmental biology, disease modeling, and therapeutic testing.^6^ Organoid models recapitulate aspects of human tissue architecture while enabling high-throughput testing of therapeutics.^7^ For example, pancreatic ductal organoids (PDOs) derived from human induced pluripotent stem cells (iPSCs) can reproduce the three-dimensional (3D) epithelial structures and key features of native ductal tissue, including luminal architecture and function.^8^ The routine practice in generating iPSC-derived organoids, including PDOs, involves the timed addition of soluble cues (*e.g.,* growth factors, pathway inhibitors) that activate or repress signaling pathways crucial at different stages of differentiation.^9^ Multi-stage differentiation protocols have been established to differentiate human iPSCs into definitive endoderm (DE), posterior foregut (PF), pancreatic progenitor (PP), and PDOs.^10^ While the DE, PF, and PP cells can be reliably differentiated in two-dimensional (2D) tissue culture plastic, all current protocols for generating PDOs rely heavily on embedding dissociated or small multicellular PP clusters in Matrigel. Despite its weak, batch-dependent, and non-tunable properties, tumor-derived Matrigel remains the *de facto* matrix in organoid research.^11^ Yet, the efficiency of generating organoids from dispersed cells embedded in Matrigel and alternative materials was generally low, such as reported in alveolar organoids (∼4%),^12^ acinar organoids (∼6%), and ductal organoids (∼2%).^13^ The low organoid formation efficiency in these examples limits the robustness of the studies and precludes high-throughput applications. Alternatively, controlled cell aggregation in spheroid-forming technologies (*e.g.*, Aggrewell^TM^) has been shown to enhance uniformity and efficiency of intestinal,^14^ neural,^15^ renal progenitor,^16^ islets,^17^ kidney,^18^ and pancreatic ductal organoids.^19^

Cell fate processes are guided by soluble morphogens, cell-cell communication, and cell-matrix interactions.^20,21^ Extracellular matrix (ECM) proteins, such as fibronectin and vitronectin, are now routinely used as 2D surface coatings to maintain and expand human iPSCs. 3D hydrogels are also increasingly used to culture or differentiate human iPSCs *in vitro,*^22–30^ including decellularized matrix (dECM) hydrogels.^31–33^ As with collagen gels and Matrigel, however, crosslinking of conventional dECM hydrogels is routinely achieved via temperature-induced hydrophobic interactions.^34^ While convenient, physical gelation often results in matrices with weak mechanical properties (shear moduli G’ < 200 Pa), which makes it challenging to induce mechanosensing in the cells. To overcome this challenge, dECM can be modified with methacrylate groups to permit chain-growth polymerization into hydrogels with higher stiffness than their thermally crosslinked counterparts.^35,36^ However, chain-growth polymerization produces high concentrations of propagating radicals even at low photoinitiator concentrations (0.05% or 1.7 mM lithium arylphosphonate, LAP).^37^ Chain-polymerization also leads to high molecular weight and heterogeneous crosslinks, producing high variation in local matrix mechanics and undesirable protein-polymer conjugates.^38,39^ Alternatively, the intrinsic tyrosine residues of the protein components of dECM have been leveraged for photocrosslinking, but this approach is limited by the scarcity of tyrosine and the need to use multiple initiators (*e.g.,* ruthenium/sodium persulfate) to initiator crosslinking.^40^

To develop a Matrigel replacement for tissue engineering and regenerative medicine applications, we recently reported a photo-clickable dECM by modifying bovine decellularized small intestinal submucosa with norbornene (*i.e.,* dSIS-NB).^41^ Different from SIS-based solid scaffolds,^42^ dSIS-NB can be crosslinked into water-swollen hydrogels via thiol-norbornene photo-click reaction. The stiffness of dSIS-NB hydrogels can be adjusted via tuning the content of thiolated macromers, such as 4-arm PEG-thiol (PEG4SH), without altering the content of cell adhesive matrices. We have also identified three major protein components in bovine dSIS-NB, including ∼60% collagen I, ∼20% collagen III, and ∼10% fibrillin I. Because collagen and fibrillin activate distinct integrins, we hypothesize that bovine dSIS-NB hydrogels could provide a highly bioactive niche to enhance stem cell differentiation efficiency. To test this hypothesis, we first generated PP cells from iPSCs in 2D culture, then aggregated them into uniformly sized PP spheroids (PPS) in Aggrewell. PPS were then encapsulated into dSIS-NB hydrogels for pancreatic ductal differentiation. We evaluated the maturation and function of PDOs, with Matrigel as a control. We also conducted single-cell RNA sequencing (scRNA-seq) to profile stage-specific cellular phenotypes in the developing PDOs.

## 2 Results

### 2.1 Differentiation of human iPSCs into PPs

Human iPSCs were first differentiated into PP cells using the 4-stage Pancreatic Progenitor Kits in 2D tissue culture plates coated with vitronectin (**Figure 1a, S1a**). After initiation of Stage 1 (S1) definitive endoderm (DE) differentiation, a substantial number of floating cells were observed, similar to that observed in routine medium exchanges. The attached cells were still highly proliferative and fully covered the surface (**Figure 1b**). Breakage of the cell monolayer was occasionally observed during S2 differentiation (**Video S1**), suggesting altered cell-cell and cell-matrix adhesion. Although a similar phenomenon has been reported using different protocols and cell lines,^43^ the molecular mechanisms remain largely unexplored. A potential mechanism underlying the weakened cell-matrix adhesion and cell shedding from the monolayer was epithelial-mesenchymal transition (EMT). Upon reaching Stage 4 (S4: pancreatic progenitors, PP), large folds of cell clusters formed from the otherwise flat monolayer (**Figure 1b**). These cells were also resistant to TrypLE dissociation, unless with a longer incubation time (over 10 minutes).

**Figure 1.**
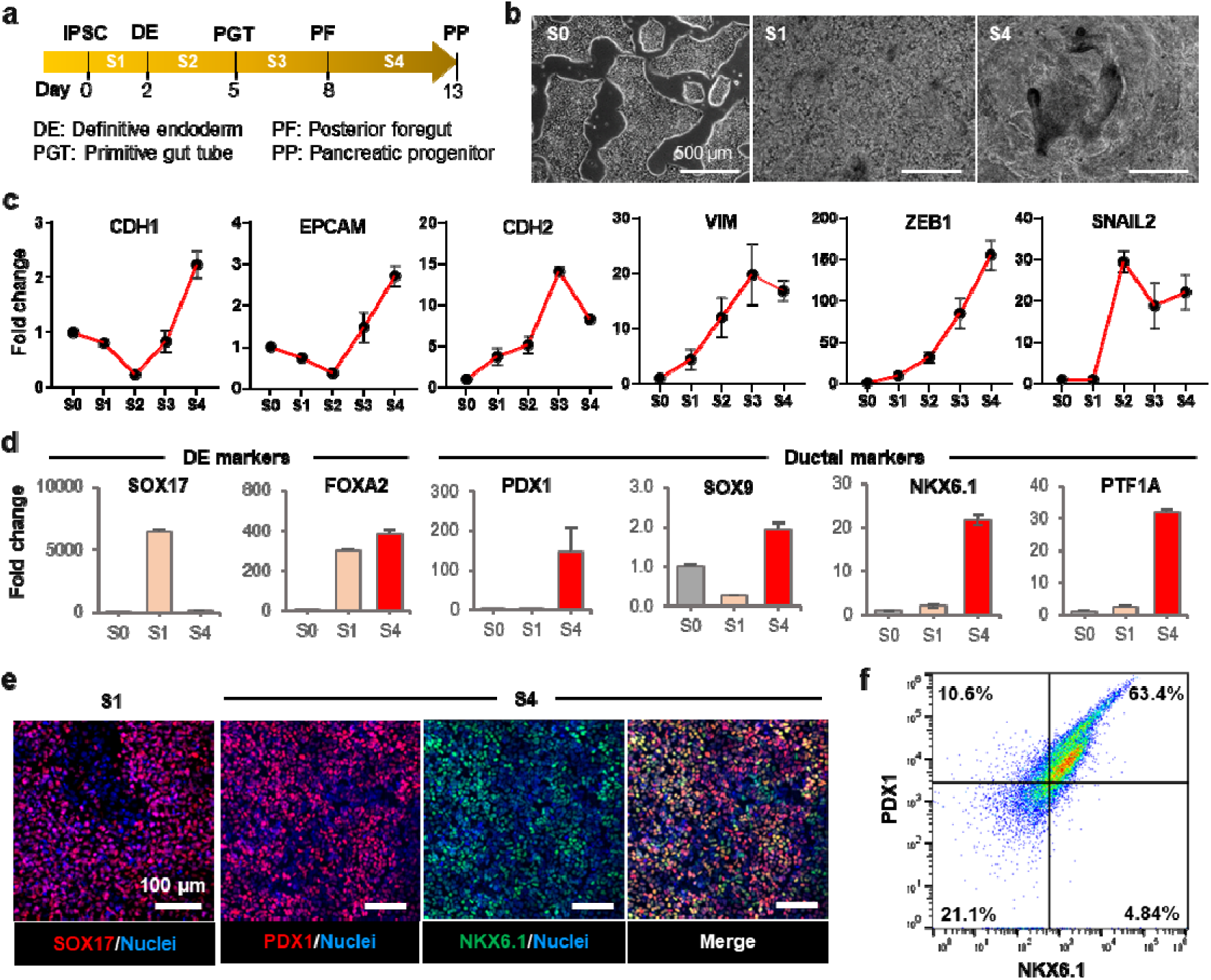
Differentiation of human iPSCs to pancreatic progenitors in 2D culture. a) Schematic of the 4-stage differentiation of iPSCs to definitive endoderm (DE), primitive gut tube (PGT), posterior foregut (PF), and pancreatic progenitors (PP). b) Representative images of morphological changes during differentiation. c) qRT-PCR analysis of key epithelial (*CDH1*, EPCAM) and mesenchymal markers (*CDH2*, *VIM*, *ZEB1*, *SNAIL2*) (n = 3; mean ± SEM). d) qRT-PCR analysis of lineage-associated markers at the indicated stages. Expression of DE markers (*SOX17*, *FOXA2*) is enriched at early stages, whereas pancreatic progenitor-associated genes (*PDX1*, *SOX9*, *NKX6.1*, *PTF1A*) are upregulated at later stages (S4) (n = 3; mean ± SEM). e) Immunofluorescence staining for SOX17 (red) at S1 and for pancreatic progenitor markers PDX1 (red), NKX6.1 (green), and nuclei (blue) at S4. f) Flow cytometric analysis of PDX1 and NKX6.1 expression at S4, with percentages indicating the proportion of cells expressing either or both markers.

To assess the potential epithelial-mesenchymal plasticity (i.e., EMT and MET) during the 4-stage PP differentiation, we performed quantitative RT-PCR (qRT-PCR) to evaluate the expression of key epithelial (*EpCAM*, *CDH1*) and mesenchymal-associated genes (*VIM*, *ZEB1, SNAIL2*) (**Figure 1c**). As iPSCs (S0) were differentiated into DE (S1) and PGT (S2) cells, expression of epithelial markers (*CDH1* and *EpCAM*) drastically decreased to only 0.1 to 0.2-fold of that in S0. At the PF (S3) and PP (S4) stages, *CDH1* and *EpCAM* expression recovered and increased up to 3-fold relative to S0. In contrast, the levels of mesenchymal markers (*CDH2*, *SNAIL2*, *VIM*, and *ZEB1*) increased progressively from S0 to S3. By S4, expression of mesenchymal markers *CDH2*, *SNAIL2*, and *VIM* decreased, suggesting a reversal toward an epithelial phenotype. These longitudinal gene expression profiles indicated a transient EMT during PP differentiation. Furthermore, DE markers *SOX17* and *FOXA2* were significantly upregulated after S1, with *SOX17* expression decreased substantially as cells transitioned to the PP stage (**Figure 1d**). Additionally, S4 cells expressed high levels of pancreatic progenitor markers *PDX1*, *SOX9*, *NKX6.1*, and *PTF1A* (**Figure 1d**). In contrast, the expression of stem cell markers *OCT4* and *NANOG* was significantly decreased at S4 (**Figure S1b)**. Gene expression results were corroborated by immunofluorescence (IF) staining (**Figure 1e**) and flow cytometry (**Figure 1f**). The PP differentiation efficiency (*i.e.,* PDX1+/NKX6.1+ cells) was above 63%, which is at the upper end compared with published values.^44^ These cells were used directly for subsequent PDO differentiation.

### 2.2. Generation of PDO in engineered dSIS-NB hydrogels

Toward replacing Matrigel for matrix-based PDO differentiation, we developed a protocol involving the generation of PP spheroids (PPS) that were subsequently encapsulated in bovine dSIS-NB hydrogels (**Figure S1a**).^41,45^ dSIS was isolated from food-grade bovine small intestine and subsequently decellularized according to the published protocol (**Figure 2a**).^41^ dSIS-NB was synthesized by reacting amine groups in dSIS with carbic anhydride in a basic (pH 8.0) aqueous buffer solution (**Figure 2b**). Hydrogel crosslinking was achieved by thiol-norbornene photo-click reaction between dSIS-NB and PEG4SH (**Figure 2c**), with gelation kinetics evaluated by *in situ* photorheometry (**Figure 2d**). Thiol-norbornene gelation of dSIS-NB/PEG4SH was fast, achieving gel points (time at which G’ surpassed G”) within 2 to 3 seconds after light was switched on (**Figure 2d**). At a fixed dSIS-NB content (1.2 wt%), adjusting PEG4SH content from 0.2 wt% to 1.2 wt% led to significantly different shear moduli (G’), ∼0.9 kPa and ∼2.5 kPa, respectively (**Figure 2e**).

**Figure 2.**
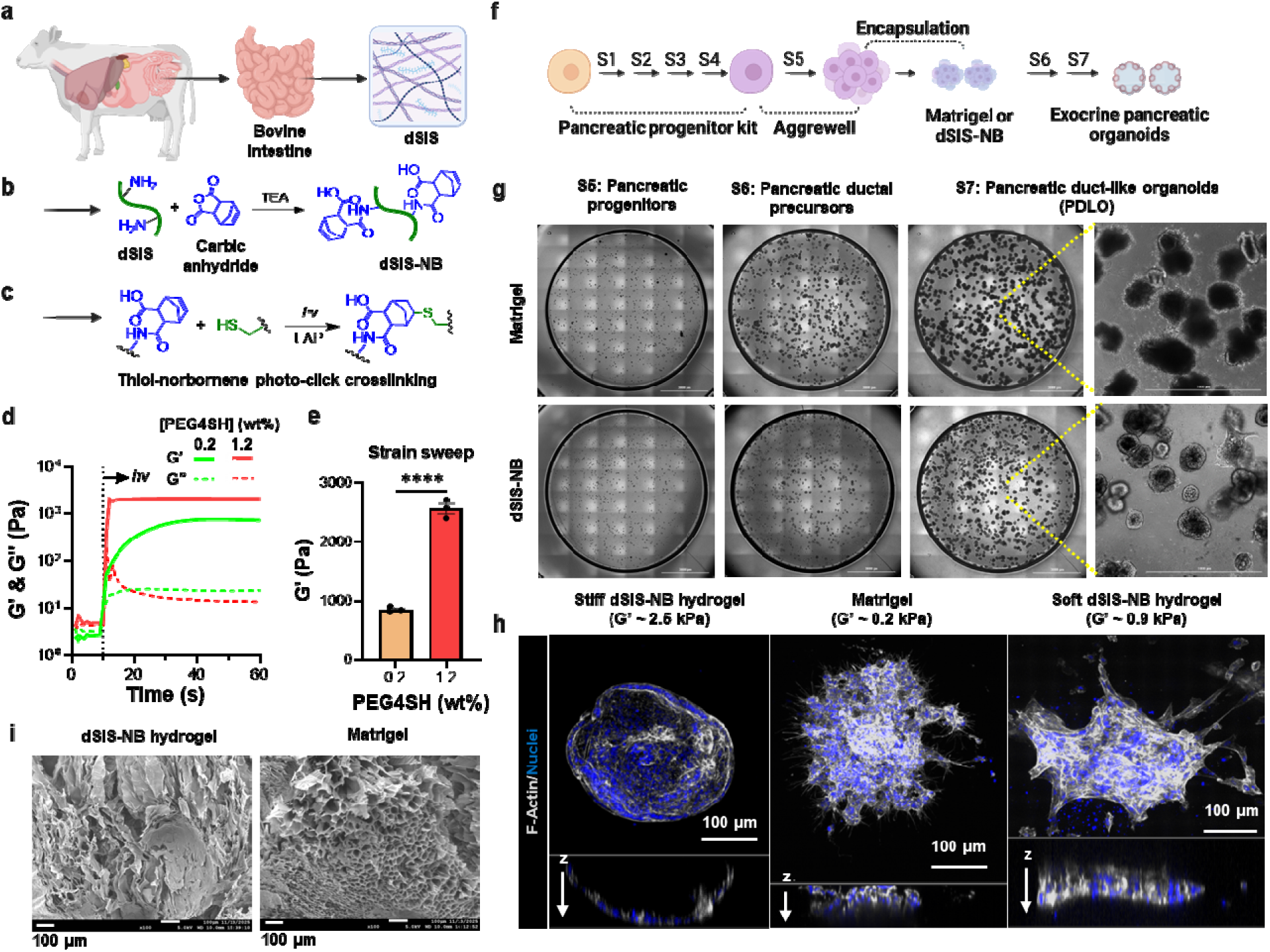
Synthesis of dSIS-NB hydrogels for differentiating iPSC-derived PPS into PDO. (a) Schematic of bovine decellularized small intestinal submucosa (dSIS) preparation (created with BioRender.com). (b) Chemical modification of dSIS to dSIS-NB via reaction with carbic anhydride. (c) Scheme of thiol-norbornene photo-click chemistry. (d) *In situ* photo-rheometry showing gelation kinetics of dSIS-NB hydrogels upon light exposure. PEG4SH (10 kDa) was used at 0.2 wt% or 1.2 wt%. Light was turned on at 10 seconds. (e) Strain sweep rheometry of dSIS-NB hydrogels crosslinked with 0.2 wt% and 1.2 wt% PEG4SH (*n* = 3; mean ± SEM; ****, *P* < 0.001, two-tailed *t*-test). (f) Schematic of PP differentiation (S1-S4), PPS aggregation (S5), and PDO differentiation (S6-S7). (g) Representative bright-field image montage of PPS encapsulated in Matrigel or dSIS-NB hydrogels during stages S5-S7. (h) Representative confocal images of F-actin staining illustrating morphological differences between spherical PDOs formed in stiffer dSIS-NB hydrogels and mesenchymal protrusions formed in Matrigel and softer dSIS-NB hydrogels. (i) Scanning electron microscopy (SEM) images of dried dSIS-NB hydrogel and Matrigel.

Photo-click dSIS-NB hydrogels were used to encapsulate uniformly sized PPS, which were formed in AggreWell™ 400 microwell plate (**Figure S1c**). The encapsulated PPS were subjected to stages 6 and 7 of PDO differentiation (**Figure 2f, S1d**), with Matrigel as a control. Regardless of the matrix used, encapsulated PPS progressively expanded in size over time (**Figure 2g, S1d, S1e**). However, PPS differentiated in Matrigel displayed disorganized morphologies with few lumens and extensive cellular invasion into the matrix (**Figure 2g**). Additionally, numerous spindle-shaped cells detached from the aggregates and migrated individually, with some invading into neighboring organoids or dividing while in motion (**Video S2**). Migrating cells exhibited spindle-like morphologies and filopodia-like extensions, characteristic of a mesenchymal phenotype. In contrast, PDO generated in the stiffer dSIS-NB hydrogels (G’ ∼2.5 kPa) remained highly viable (**Figure S1f**) and formed compact cystic structures with clear central lumens and epithelial boundaries (**Figure 2g**, **Video S3**). Both dSIS-NB hydrogels and Matrigel supported a high efficiency of outgrowth structure formation (**Figure S1g)**, suggesting that both matrices provide a permissive microenvironment for structure expansion. Confocal imaging of F-actin confirmed well-defined luminal organization in the PDOs formed in stiffer dSIS-NB hydrogels (**Figure 2h, Video S4, right**). On the other hand, PPS maintained and differentiated in both softer dSIS-NB gels (G’ ∼0.9 kPa) and Matrigel (G’ ∼ 0.2 kPa), which displayed disrupted junctions (**Figures 2h**, **S2**, **Video S4, left**). Given its superior structural stability and support of organoid integrity, the stiffer dSIS-NB hydrogel was chosen for all subsequent experiments. Matrigel was used as the benchmark as it is the current ‘gold-standard’ for organoid research.

Because dSIS-NB hydrogels and Matrigel were crosslinked via distinct gelation mechanisms (i.e., photocrosslinking vs. thermal gelation), the microstructure of the hydrogel network may be a confounding factor contributing to the observed morphological differences. Hence, the microstructures of dSIS-NB hydrogel and Matrigel were evaluated by scanning electron microscopy (SEM). SEM images of dried substrates of dSIS-NB and Matrigel revealed that both materials exhibited a ‘pseudo-porous’ (due to the vacuum drying process) and non-fibrous structure (**Figure 2i**). While dSIS-NB is composed of fibril-forming collagen and fibrillin, we have shown that chemical crosslinking of dSIS-NB hydrogels diminished the fibrous structure.^41^ On the other hand, basement membrane-derived Matrigel is rich in collagen IV and laminin, which assemble into a felt-like meshwork. Nonetheless, the two matrices exhibited indistinguishable microstructures, albeit crosslinked by different mechanisms and with different stiffnesses, thus ruling out microstructure as an underlying cause of the observed morphological variation. To assess the robustness of photo-click dSIS-NB hydrogels in organoid differentiation, a different human iPSC line (WTC-mEGFP-CTNNB1-cl67) was used to derive PDOs. Cystic morphology and epithelial organization were again observed in photo-click dSIS-NB hydrogels. In contrast, chestnut-like structures were observed in Matrigel culture (**Figure S3**). Collectively, these results demonstrate that photo-click dSIS-NB hydrogels enable robust, highly efficient formation of PDOs.

### 2.3 Characterization of PDO Maturation

To evaluate how different matrices influence epithelial organization during PDO maturation, we compared organoids cultured in dSIS-NB hydrogels with those in Matrigel. IF images of PDO formed in dSIS-NB hydrogels exhibited a continuous, lumen-enclosing epithelial layer with uniform E-cadherin and SOX9 expression, whereas Matrigel-cultured PPS displayed disorganized morphology and fragmented epithelial domains (**Figure 3a**). dSIS-NB-derived PDOs also showed enriched EpCAM expression with limited vimentin expression, indicative of a predominantly epithelial cell composition. In contrast, PPS in Matrigel expressed notable mesenchymal marker vimentin (**Figure 3b**). Furthermore, only PDO formed in dSIS-NB gels were properly polarized, with apical surface stained positive for ZO-1 and basal lateral surface stained positive for EpCAM (**Figure 3c**). Next, ductal markers KRT7, KRT19, and PDX1 were highly expressed in dSIS-NB-derived PDOs, but not in Matrigel control (**Figure 3d, e**). In addition, a subset of dSIS-NB-derived PDOs underwent morphological elongation at the end of S7, giving rise to non-spherical organoids (**Figure 3f**). The maturation of dSIS-NB-derived PDOs was further confirmed by the upregulation of *CDH1* and *EpCAM*, and downregulation of *VIM* and *ZEB1* at the mRNA level (**Figure 3g**). The mRNA expression of ductal marker genes (*KRT19, KRT7, SOX9*, and *PDX1*) was upregulated in dSIS-NB-derived PDOs relative to Matrigel control (**Figure 3h**). mRNA expression of acinar (*CEL*, *GATA4*) and endocrine marker genes (*INS*, *NKX2.2*) remained low in both matrices (**Figure S4**), indicating that differentiation was specifically biased toward ductal identity. Moreover, the expression levels of endothelial markers (*CD31*, *CDH5*) were high at S4, but reduced when reaching S7 (**Figure S4**), indicating minimal off-target differentiation. Together, these data reaffirm that photo-click dSIS-NB hydrogels supported robust PDO maturation.

**Figure 3.**
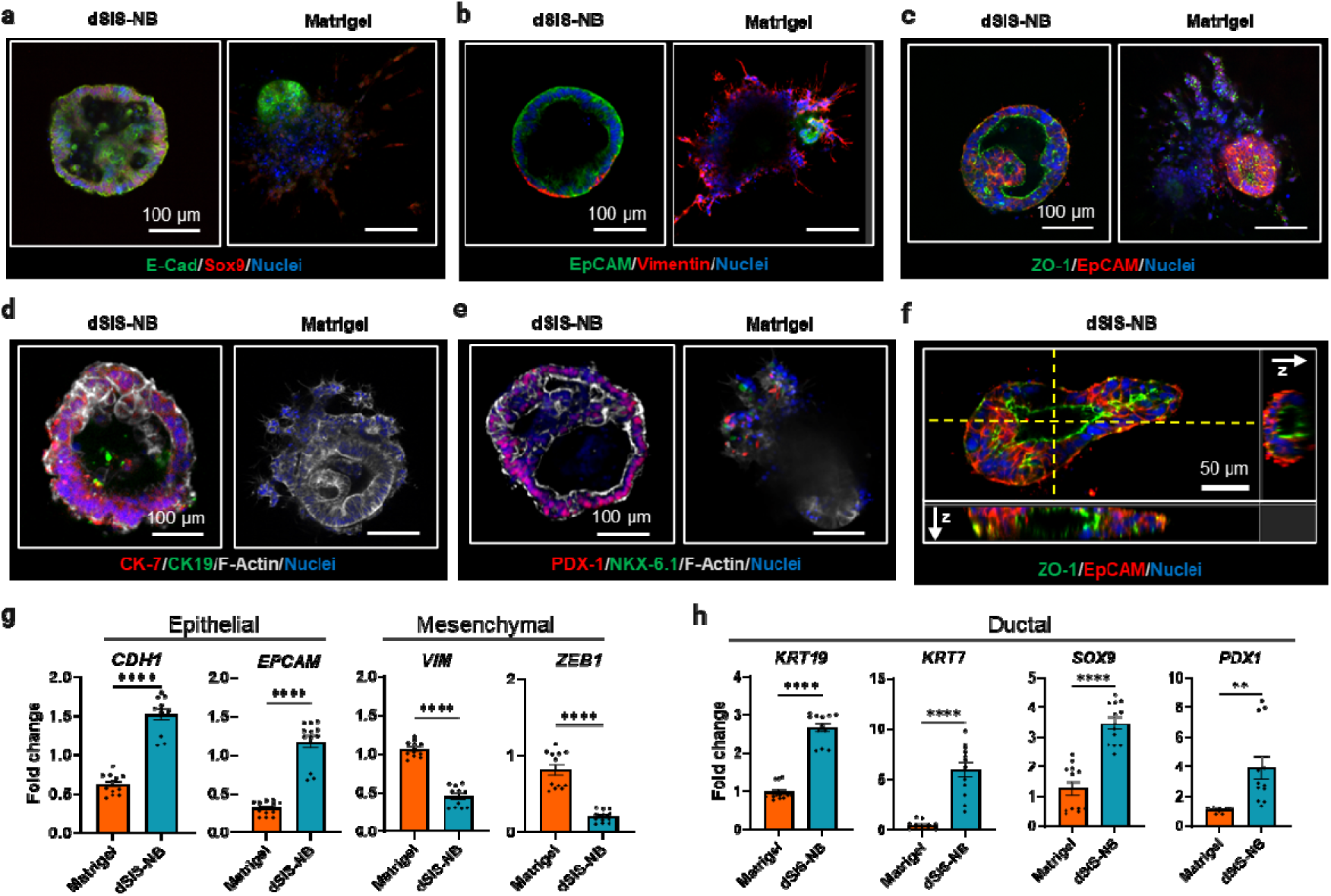
dSIS-NB hydrogels support epithelial organization and ductal marke expression in PDOs. a-e) Representative immunofluorescence staining images of PDOs cultured in dSIS-NB hydrogels or Matrigel: a) E-cadherin (green) and SOX9 (red). b) EpCAM (green) and vimentin (red). c) ZO-1 (green) and EpCAM (red). d) CK7 (red), CK19 (green), and F-actin (white). e) PDX1 (red), NKX6.1 (green), and F-actin (white). f) Orthogonal (x-z and y-z) confocal views of a dSIS-NB-cultured PDO stained for ZO-1 (green), EpCAM (red), and nuclei (blue), illustrating elongated epithelial polarization and lumen formation. g) qRT-PCR analysis of epithelial (*CDH1*, *EPCAM*) and mesenchymal-associated (*VIM*, *ZEB1*) gene expression. h) qRT-PCR analysis of ductal-associated gene expression (*KRT19*, *KRT7*, *SOX9*, *PDX1*). Fold change relative to PPS; n = 4 independent batches; data are presented as mean ± SEM; **p < 0.01, ****p < 0.0001.

### 2.4 Single-cell RNA sequencing (scRNA-seq) verifies PDO maturation in dSIS-NB hydrogel

To assess the impact of different matrices on PDO maturation, we performed scRNA-seq on S7 cells derived from either Matrigel or dSIS-NB hydrogels. Global differential expression analysis revealed extensive transcriptional divergence. Among 26,534 genes detected, 601 were significantly different in expression levels, with 458 upregulated and 143 downregulated in dSIS-NB-derived PDOs relative to Matrigel (**Figure 4a, Supplementary Table S1**). Functional enrichment analysis highlighted a pronounced metabolic and developmental shift between the two matrices. Specifically, dSIS-NB-derived PDOs were significantly enriched for nuclear-encoded oxidative phosphorylation (OXPHOS) and mitochondrial biogenesis pathways (**Figure 4b**, **left**). This was evidenced by the coordinated upregulation of key mitochondrial-encoded genes (*e.g., MT-ND1-5*, *MT-CO2/3*, and *MT-ATP6/8*) and nuclear-encoded OXPHOS components (*e.g., NDUFA* and *NDUFB* families, *COX6C*, and *ATP5* family) (**Supplementary Table S1**). In contrast, Matrigel-grown organoids were enriched for canonical glycolysis programs (**Figure 4b**, right), characterized by higher expression of glycolytic enzymes (*LDHA*, *ENO1*, *PKM*, *ALDOA*, *TPI1*) and the glucose transporter *SLC2A1* (*GLUT1*) (**Supplementary Table S1**). Consistently, Gene ontology (GO) biological process analysis confirmed strong representation of terms associated with mitochondrial ATP synthesis and oxidative phosphorylation in the dSIS-NB group, whereas Matrigel-upregulated genes reflected a progenitor-like profile enriched for ribosome biogenesis, cell-cycle regulation, and glycolysis (**Figure 4c, Supplementary Table S2**). The pronounced enrichments of OXPHOS and mitochondrial biogenesis pathways in dSIS-NB-derived PDOs are consistent with previous reports, where the hallmark of differentiation and functional maturation is accompanied by a metabolic shift from glycolytic metabolism in the pluripotent cells to oxidative phosphorylation in the differentiated cells.^46–49^

**Figure 4.**
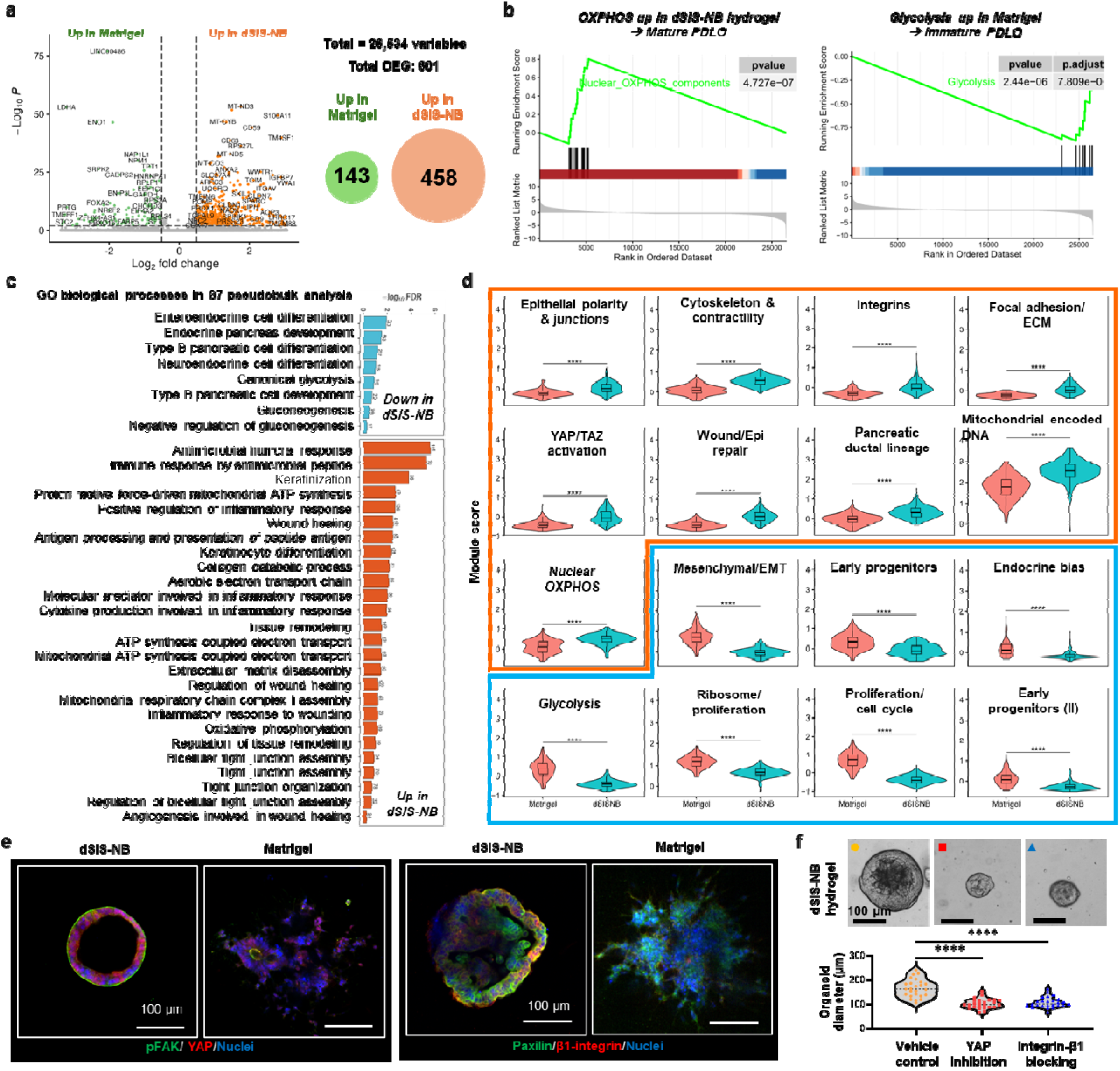
Transcriptomic profiling reveals metabolic reprogramming and enhanced mechanotransduction in dSIS-NB-derived PDO. a) Volcano plot showing differentiall expressed genes (DEGs) between S7 PDOs cultured in dSIS-NB hydrogels versus Matrigel. Dashed lines indicate significance thresholds (log₂ fold change > |0.25|, adjusted p < 0.05). Venn diagram illustrating the number of significantly upregulated genes in Matrigel (143) and dSIS-NB (458). b) Gene Set Enrichment Analysis (GSEA) plots showing a significant enrichment of Oxidative Phosphorylation (OXPHOS) signatures in dSIS-NB-derived PDOs (left) versus Glycolysis signatures in Matrigel-derived cells (right). c) Top enriched Gene Ontology (GO) Biological Processes based on pseudobulk DEGs. The dSIS-NB group (orange bars) is enriched for terms related to wound healing, mitochondrial ATP synthesis, and tight junction assembly, whereas the Matrigel group (blue bars) is enriched for glycolysis and endocrine differentiation terms. d) Violin plots of module scores for manually curated gene sets. The orange box highlights modules upregulated in dSIS-NB (including Epithelial polarity, Integrins, YAP/TAZ activation, and Mitochondrial OXPHOS), while the blue box highlights modules upregulated in Matrigel (Mesenchymal/EMT, Glycolysis, and Proliferation). Statistical significance was determined by Wilcoxon Rank Sum test (**** p < 0.0001). e) Representative IF staining of PDOs in dSIS-NB gel and Matrigel, showing expression of Paxilin, β1-integrin, pFAK, and YAP. f) Functional perturbation of metabolic and mechanotransduction pathways impairs PDO growth. PDOs cultured in dSIS-NB hydrogels were treated with vehicle control (●: 0.1% DMSO), YAP inhibitor (▪: 5 nM verteporfin), or integrin-β1 blocking (▴: 2.5 µg/mL blocking antibody) during S7 differentiation. Organoid diameters were quantified at the end of differentiation.

Beyond metabolism, module-score analysis of curated signatures revealed distinct structural trajectories (**Figure 4d**). dSIS-NB-derived PDOs exhibited coordinated upregulation of programs involved in epithelial polarity, barrier assembly, and keratinization, alongside signatures of wound healing and ECM organization. Conversely, Matrigel-cultured organoids scored higher for mesenchymal/EMT modules and early-progenitor programs, suggesting the maintenance of a less differentiated phenotype. The remodeling signature in dSIS-NB hydrogel appears to be mechanically driven, as modules associated with actomyosin contractility, integrins, focal adhesion, and YAP/TAZ-responsive transcription were robustly enriched. We also found that dSIS-NB hydrogels significantly upregulated a broad range of integrin subunits, including *ITGB1 (*β*1), ITGB6 (*β*6) ITGB8 (*β*8), ITGAV (*α*v), ITGA2 (*α*2), and ITGA3 (*α*3)* (**Supplementary Table S1**). To determine how ECM cues govern epithelial growth and function during late-stage PDO maturation, we first compared YAP activity and integrin signaling in S7 PDOs cultured in dSIS-NB hydrogel or Matrigel. Immunofluorescence staining results revealed enhanced nuclear localization of YAP and increased integrin β1 organization at the epithelial-matrix interface, as well as expression of Paxilin and pFAK in PDOs cultured in dSIS-NB relative to Matrigel (**Figure 4e**). To interrogate the contribution of these pathways to PDO maturation, we added an integrin-β1 blocking antibody (2.5 µg/mL) or a YAP inhibitor (5 nM verteporfin) during S7 differentiation. Under these conditions, PDOs exhibited a marked reduction in organoid size compared with untreated controls, indicating impaired epithelial expansion during ductal maturation (**Figure 4f**).

### 2.5 Forskolin-Induced Swelling Confirms Functional CFTR Activity

Next, maturation of PDOs was verified by detecting the expression of Cystic Fibrosis Transmembrane Conductance Regulator (CFTR), a chloride and bicarbonate channel critical for pancreatic ductal secretion (**Figure 5a, b**). Furthermore, only PDOs generated in photo-click dSIS-NB hydrogels were highly responsive to forskolin-induced swelling (FIS) (**Figure 5c, d, video S5**). Forskolin-induced luminal expansion is CFTR-dependent and the loss of CFTR expression leads to non-functional ductal tissues.^50^ To further confirm that the swelling observed in dSIS-NB-derived PDOs was specifically mediated by CFTR activity, we combined forskolin with CFTR inhibitor-172 (50 µM) (**Figure S5)**. When the inhibitor was added for 2 h prior to forskolin treatment, the swelling response was completely blocked, with lumen sizes comparable to untreated controls. Similarly, when CFTR inhibitor-172 was applied after 24 h forskolin stimulation, the enlarged lumens were prevented from further expansion, with some organoids showing a slight reduction in size (**Figure S5**). Time-lapse imaging demonstrated that swelling peaked at approximately 36 h after forskolin stimulation without CFTR inhibitor-172 (positive control) and regressed thereafter. In contrast, PDOs exposed to DMSO (untreated control) continued to grow at a normal rate without acute swelling, confirming that the effect was specific to forskolin-induced CFTR activation (**Figure S5**).

**Figure 5.**
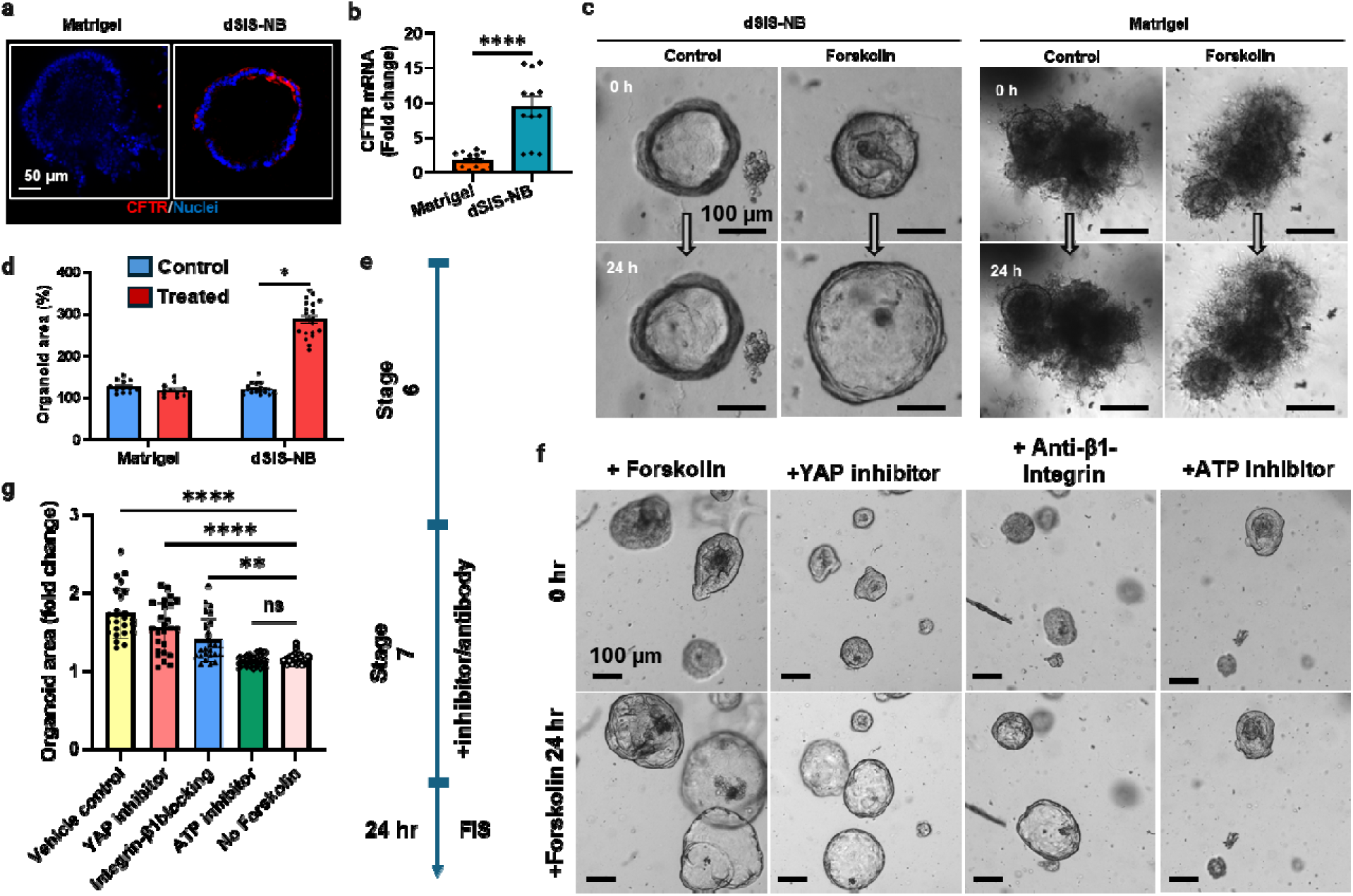
Forskolin-induced swelling reveals enhanced CFTR-dependent ductal function in PDOs cultured in dSIS-NB hydrogels. a) Representative immunofluorescence staining of CFTR in PDOs cultured in Matrigel or dSIS-NB hydrogels. b) qRT-PCR detection of CFTR mRNA expression levels in PDOs cultured in Matrigel or dSIS-NB hydrogels. 1-fold: Relative to PP controls. ****P < 0.0001. c) Representative bright-field images of forskolin-induced swelling (FIS) assays. PDOs cultured in dSIS-NB hydrogels or Matrigel were imaged pre-treatment (0 h) and after 24 h of forskolin treatment. d) Quantification of organoid area before and after forskolin treatment (*p < 0.05). e) Experimental timeline and f) representative images showing the effects of pathway perturbation during stage 7 on FIS responses post differentiation. PDOs were treated with vehicle control, YAP inhibitor (5 nM verteporfin), 2.5 µg/mL β1-integrin-blocking antibody, or mitochondrial ATP synthase inhibitor (5 µM oligomycin), followed by forskolin stimulation for 24 h. g) Quantification of organoid area following forskolin treatment, expressed as fold change relative to pre-treatment area.

We then compared FIS of PDOs cultured in dSIS-NB under control conditions or following pathway perturbation, such as integrin, YAP or mitochondrial ATP at the end of S7 (**Figure 5e**). PDOs treated with 2.5 µg/mL integrin-β1 blocking antibody or 5 nM verteporfin retained the ability to swell in response to forskolin; however, the magnitude of swelling was reduced relative to vehicle controls (**Figure 5f, 5g**). This partial response suggests that while integrin/YAP signaling is not absolutely required for CFTR activation, it is necessary for achieving full ductal functional competence. In contrast, PDOs treated with 5 µM oligomycin, a mitochondrial ATP synthase inhibitor, exhibited little to no swelling response following forskolin stimulation. The near-complete loss of CFTR-dependent swelling under mitochondrial inhibition indicates a critical requirement for mitochondrial ATP production in supporting ductal ion transport and fluid secretion. Together, these data demonstrate that integrin-β1/YAP signaling and mitochondrial metabolism play distinct but complementary roles in PDO maturation: integrin-YAP signaling primarily governs epithelial organization and growth, whereas mitochondrial oxidative phosphorylation is essential for executing CFTR-dependent ductal function.

### 2.6 Tracking pancreatic ductal development via scRNA-seq

Encouraged by the matured and functional PDO generated in dSIS-NB hydrogels, we sought to resolve the cellular heterogeneity and developmental trajectories underlying this maturation. We performed scRNA-seq on cells collected from dSIS-NB hydrogels at three key differentiation stages: S5 (spheroid aggregation), S6 (ductal specification), and S7 (ductal maturation). The dataset comprised well-resolved cellular clusters, as shown in the uniform manifold approximation and projection (UMAP) (**Figure 6a, b**). To achieve robust cell-type annotation, we integrated hierarchical clustering (**Figure 6c**), differential gene-expression analysis (**Figure 6d**), and canonical marker profiling (Figure 6e; **S6**). In addition, pseudotime trajectory analysis was employed to reconstruct the lineage progression from progenitors to differentiated cells (**Figure 6f**).

**Figure 6.**
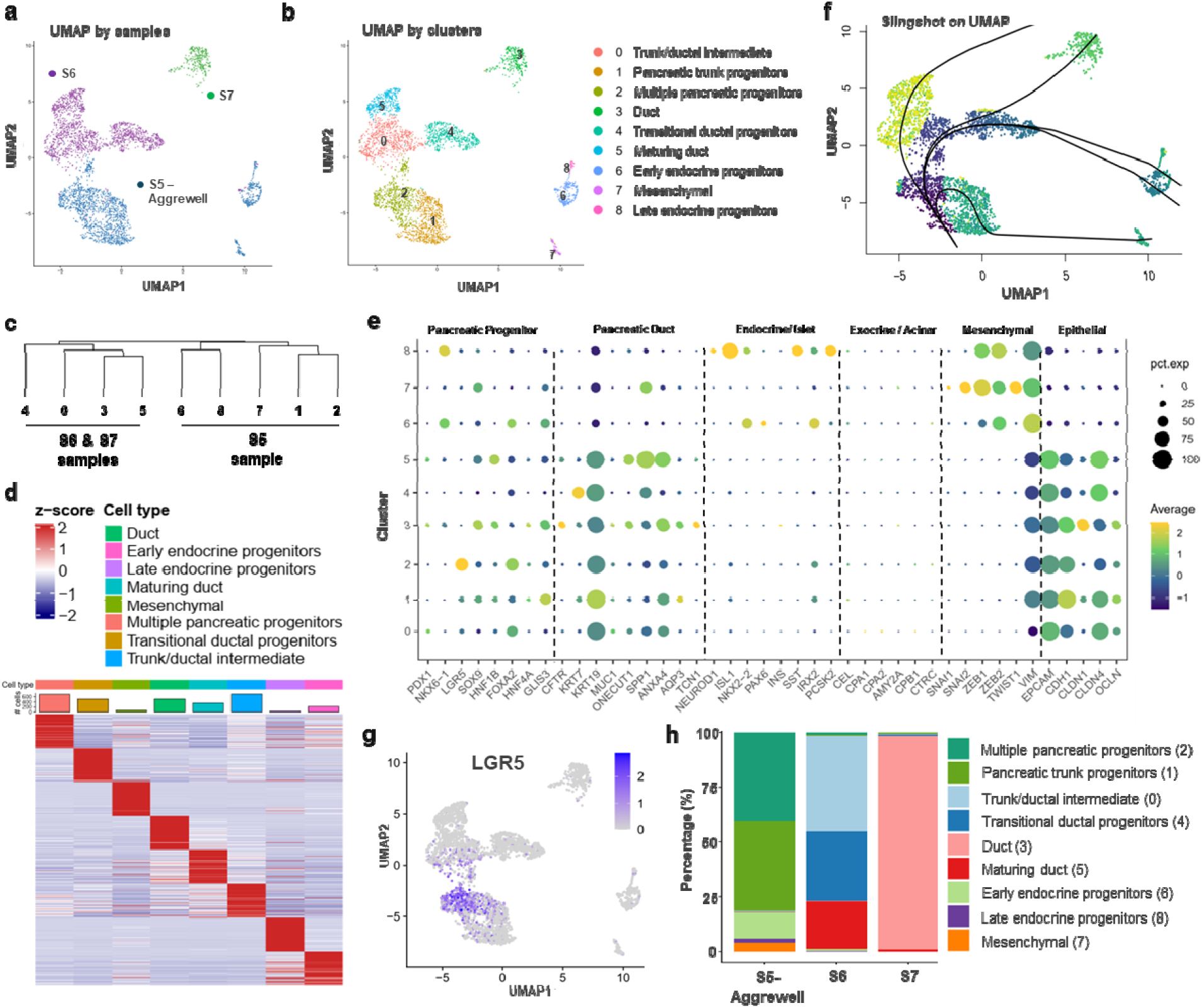
Single-cell transcriptomic analysis and cell-type annotation of PDOs cultured in dSIS-NB. a) UMAP of all single cells passing quality control colored according to the developmental stage of each sample (S5, S6, and S7), illustrating the progressive transition from pancreatic progenitors to mature ductal phenotypes in dSIS-NB hydrogel. b) UMAP of all single cells passing quality control, showing well-resolved clusters corresponding to distinct cellular populations across differentiation stages. Nine clusters were annotated according to their gene expression patterns. c) Hierarchical clustering dendrogram depicting transcriptional relationships among identified clusters. d) Heatmap displaying the scaled expression (z-score) of the top 100 differentially expressed genes for each cluster. e) Dot plot characterizing cluster identities showing the expression of representative marker genes across the nine identified clusters. Rows indicate clusters and columns indicate genes. Dot size represents the percentage of cells expressing the gene, and color intensity indicates average expression level. f) Pseudotime trajectory analysis revealing the direction and magnitude of differentiation transitions among cell populations, consistent with a continuous epithelial ductal maturation process. g) UMAP visualization highlighting cells with detectable LGR5 expression. h) Stacked bar plot quantifying the dynamic shifts in cell-type proportions across differentiation stages, highlighting the progressive enrichment of ductal cells.

At S5, five transcriptionally distinct clusters were identified: including pancreatic trunk progenitors (cluster 1), multipotent pancreatic progenitors (cluster 2), early endocrine progenitors (cluster 6), mesenchymal cells (cluster 7), and late endocrine progenitors (cluster 8) (**Fig. 6a, b**). Clusters 1 and 2 expressed canonical pancreatic progenitor markers, such as *HNF1B*, *PDX1*, *NKX6-1* and *SOX9* (**Figure 6e, S6**). Notably, Cluster 2 was distinguished by the enrichment of *LGR5*, a marker of tripotent pancreatic stem/progenitor cells (**Figure 6g**), positioning it as the putative root of the developmental hierarchy. In contrast, Cluster 1 exhibited a ductal bias characterized by high *KRT19* expression. Clusters 6 and 8 expressed lower levels of ductal progenitor markers (*HNF1B* and *ANXA4*) but high levels of genes associated with early endocrine progenitors (*NKX2.2, PAX6* in cluster 6) and late endocrine progenitors (*NEUROD1*, *PCSK2, SST,* and *ISL1* in cluster 8). The cluster dendrogram (**Figure 6c**) and heatmap of the top 100 expressed genes (**Figure 6d**) showed a close relationship between clusters 1 and 2, as well as between clusters 6 and 8, suggesting a shared core gene-expression program among these clusters. Finally, cluster 7, a distantly separated branch with a unique gene expression pattern, was assigned to mesenchymal cells due to its high expression of *SNAI2*, *ZEB1* and *TWIST1* (**Figure 6e**).

At S6, transcriptional profiling resolved three major subpopulations corresponding to progressive ductal differentiation: trunk/ductal intermediate progenitors (cluster 0), transitional ductal progenitors (cluster 4), and maturing ductal cells (cluster 5). Cluster 0 retained progenitor features but displayed elevated *HES1* and *ANXA4*. Cluster 4 had reduced progenitor gene expression and higher ductal marker *KRT7*. Cluster 5 harbored cells with upregulated *SPP1,* an adult human pancreatic ductal marker (**Figure 6e, S6**). This progression was visualized by pseudotime modeling (DiffusionMaps and Slingshot), which inferred a branching trajectory originating from the LGR5⁺ progenitors (Cluster 2). This trajectory bifurcated into ductal (leading to Cluster 3 at S7) and endocrine (Clusters 6 and 8) lineages, with the dSIS-NB microenvironment efficiently supporting the progression along the ductal branch (**Figure 6f**).

By S7, cells consolidated into a predominant ductal population (cluster 3) characterized by high expression of *SPP1*, *ANXA4*, *SOX9,* and *CFTR*. Moreover, the suppression of acinar markers (*CEL*, *CPA1*/*2*, *AMY2A*, *CTRC*) further supports the conclusion that the differentiation protocol favored the ductal fate. Based on these combined analyses, we identified a clear developmental continuum from PP to PDO phenotypes, with cell populations shifted from intermediate progenitor-like clusters to fully differentiated ductal cells. By stage S7, approximately 97% of cells were annotated as ductal cells, with the remaining ∼3% classified as intermediate ductal subtypes (**Figure 6h**). This progressive consolidation of cell identity highlights the efficiency and fidelity of the differentiation process within the photo-click dSIS-NB hydrogels. Collectively, these data therefore support a model in which stiffer photo-click dSIS-NB hydrogels not only accelerate progenitor-to-ductal differentiation but also restrain alternate lineage fates, yielding a streamlined ductal epithelial population for downstream functional maturation.

## 3. Discussion

dECM hydrogels have emerged as a promising alternative to Matrigel for 3D cell culture owing to their non-tumor origin and tissue-specific composition.^51^ However, like Matrigel, conventional dECM hydrogels suffer from some of the same disadvantages of Matrigel, namely weak mechanics and lack of tunability in crosslinking. To this end, we have developed dSIS-NB, a photo-clickable dECM, that can be photocrosslinked into hydrogels with tunable mechanical properties while preserving the bioactivity of the native matrix. In this work, we leveraged dSIS-NB hydrogels to establish a Matrigel-free PDO differentiation protocol. We showed that stiffer (G’ ∼ 2.5 kPa) dSIS-NB hydrogels supported stepwise differentiation of PPS into functional PDO. Different from Matrigel where the major protein components are collagen IV and laminin, bovine dSIS is enriched with collagen I (∼60%), collagen III (∼20%), collagen V (∼3%), and fibrillin I (∼10%).^45,52^ Native collagens form a triple-helical structure and primarily bind to integrins α1β1, α2β1, α10β1, and α11β1 through a distinct recognition motif (*e.g.,* GFOGER sequence),^53,54^ whereas fibrillin activates integrins α5β1, α5β6, α8β1, and αvβ3 through its RGD motif.^55–58^ Activation of both integrins αvβ3 and α5β1 is critical for pancreatic organogenesis.^59,60^ While Matrigel can also activate integrin αvβ3, it also inhibits integrin α5β1 via laminin.^61^ These integrins are known to mediate focal adhesion formation and activate YAP/TAZ signaling to regulate epithelial morphogenesis,^62^ resulting in higher expression of several YAP target genes (*e.g., ANKRD1*, *ANXA1*, *WWTR1*, *ANKRD1*, *CCN1*/*CCN2* family. **Figure 4d**, **Supplementary Table S1**). These results align with current models of integrin-mediated matrix-guided morphogenesis.^63^ Coincidentally, our scRNA-seq data revealed upregulation of *ITGA2*, *ITGA3*, *ITGAV*, *ITGB1*, *ITGB6*, and *ITGB8* in PDOs generated in stiffer dSIS-NB hydrogels (**Supplementary Table S1**), suggesting a critical role of collagen I, III, V, and fibrillin I in promoting PDO maturation. Integrin-mediated adhesion, particularly on stiffer substrates, induces focal adhesion maturation.^64^ The outside-in mechanical transduction activates focal adhesion kinase (FAK) and Rho/ROCK signaling, leading to stress fiber formation and eventually nuclear translocation of the transcriptional co-activators YAP/TAZ.^65^ In addition to stiffness, matrix viscoelasticity also impacts organoid growth and morphogenesis via the integrin-YAP signaling axis.^66^ ^67^ In the pancreas, collagen I, IV, V, VI, and other ECM ligands engage integrins on pancreatic cells and influence their survival, differentiation, and function.^68^ Activation of integrins has also been shown to direct stem cell fate and organoid formation.^69^ Furthermore, YAP/TAZ nuclear activation in response to stiffer or more adhesive ECM is well documented.^70^

In addition to notable integrin activation, bovine dSIS-NB hydrogels also induced higher OXPHOS and lower glycolysis gene modules in PDOs than those formed in Matrigel. Studies have shown that metabolic reprogramming is frequently a downstream consequence of adhesion and tension-driven cell states. In this light, the up-regulation of OXPHOS-related genes in PDOs generated from dSIS-NB gels is consistent with the metabolic shifting from a glycolytic, progenitor-like state (which is typical of Matrigel culture) toward a more mitochondrial catabolic activity, a polarized epithelial phenotype as shown in hepatocyte,^71^ kidney,^72^ osteogenic^73^, neurons,^74^ and β cell differentiation^75^. From a functional perspective, the enhanced CFTR expression enabled forskolin-induced fluid transport and progressive swelling of PDOs in dSIS-NB hydrogels, but not in Matrigel (**Figure 3g, 3h**). While the outside-in signaling pathways leading to the maturation of PDO have been established, few, if any, engineered matrices permit the induction of these pathways for developing pancreatic ductal organoids.

Our data suggest that during differentiation from human iPSCs to PDO, cells undergo a transient EMT/MET. A peak in EMT-like gene expression was detected at early stages (S2-S3), followed by MET-like re-epithelialization in later stages (S4-S7). Such biphasic EMT/MET transitions are deeply embedded in embryonic development and organogenesis yet have seldom been described in the context of *in vitro* pancreatic lineage specification. The observed temporal transcriptional patterns of EMT regulators may facilitate progenitor rearrangement, detachment, and morphological plasticity required prior to recommitting to an epithelial ductal program. For example, EMT and MET transitions are known to underpin organ morphogenesis and branching. Although much of the literature focuses on cancer research (*e.g.,* EMT in pancreatic cancer)^76^ the developmental studies are increasingly recognized, albeit understudied. From scRNA-seq results (**Figure 5e**), we also found that Cluster 2 exhibited robust expression of *LGR5*, a hallmark of a self-renewing stem cell population previously identified in the human 14 to 15 gestation weeks fetal pancreas.^77^ These *LGR5*⁺ cells represent a tripotent progenitor state, capable of generating acinar, ductal, and endocrine lineages, as demonstrated by single-cell-derived pancreatic organoid assays. Their presence in our iPSC-PPS provides the first direct evidence that LGR5⁺ fetal-like progenitor populations can be reconstituted *in vitro* during human pancreatic differentiation. The abundance of *LGR5*⁺ progenitors at S5 declined by S6 and S7, consistent with the temporal restriction of *LGR5* expression observed during human pancreas development. Trajectory analysis using *DiffusionMaps* and *Slingshot* revealed that differentiation proceeded from the LGR5⁺ progenitor pool (cluster 2) through transitional ductal intermediates (cluster 4) toward mature ductal cells (cluster 3). In contrast, clusters 6 and 8 diverged toward endocrine fates. This bifurcated trajectory highlights the central role of LGR5⁺ cells as a progenitor node from which both ductal and endocrine lineages arise, consistent with *in vivo* human fetal pancreatic hierarchies. Together, these results provide compelling evidence that iPSC-derived pancreatic organoids can recapitulate fetal progenitor hierarchies, including the *LGR5*⁺ multipotent stem cell state.

Although this study demonstrates that a photo-click dSIS-NB hydrogel supports reproducible morphogenesis and enhanced epithelial maturation in PDOs, several limitations should be acknowledged. First, the current model is specifically optimized for ductal differentiation and does not incorporate the full cellular diversity of the pancreas, including endocrine and acinar lineages. As a result, the generated PDOs do not fully recapitulate the multicellular complexity of native pancreatic tissue. Future studies will focus on enriching and isolating defined progenitor populations, such as LGR5⁺ cells at Stages 4 and 5, and directing their differentiation toward multiple pancreatic lineages within a single organoid platform to establish a more comprehensive pancreatic organoid model. Second, although our perturbation experiments indicate that integrin–β1/YAP signaling and mitochondrial metabolism contribute to PDO maturation, the precise molecular mechanisms linking ECM composition and mechanical cues to epithelial organization and ductal function remain incompletely defined. Future studies combining genetic perturbation, controlled modulation of matrix properties, and metabolic flux analyses will be required to establish these relationships. Such efforts will advance the development of physiologically faithful organoid systems for the study of human pancreatic biology and disease.

In conclusion, our data strengthen the concept that scaffold biochemistry and mechanics jointly contribute to organoid morphogenesis. The performance of photo-clickable dSIS-NB hydrogel underscores the translational potential as a physiologically relevant ECM platform for functional organoid generation. Looking ahead, we envision that the dSIS-NB hydrogel platform will serve not only for pancreatic ductal organoid biology but also for other epithelial organoid systems in which mechanobiology and metabolic maturation are critical.

## 4. Experimental Section

### 4.1 Human induced pluripotent stem cell (iPSC) culture and expansion

Cellartis hiPSC12 cell line (ChiPSC12, Takara) were cultured on vitronectin (VTN)-coated tissue plates in Essential 8^TM^ (E8, Gibco) medium followed manufacturer’s protocol. The cells were passaged by using TrypLE^TM^ Select dissociation reagent (Gibco) when the confluence atL∼L70-80%. Cells were reseeded on VTN-coated tissue plates in E8 supplemented with 10 µM ROCK inhibitor Y-27632 (Selleck) for the first 24 hours (h). Daily complete media exchanges were done withinL±L2 h of the previous fed time. hiPSCs were passaged at least once after thawing and prior to commencing differentiation and the number of passages used for all experiments was from 35 to 40.

### 4.2 Pancreatic ductal-like organoid differentiation from iPSCs

To initiate differentiation, confluent iPSCs were dissociated into single-cell suspensions using TrypLE, counted and seeded at 2.5L×L10^5^ cells/cm^2^ into vitronectin-coated well plates in E8 medium supplemented with 10 μM Y27632 (Selleck). Differentiation was initiated when cells reached 80-90 confluence. The differentiation medium is changed daily according to the stepwise differentiation protocol of the STEMdiff™ Pancreatic Progenitor Kit (STEMCELL, Cat#05120). The differentiation process, including the media and factors used at each stage, is illustrated in **Supplemental Figure S1a**. Following continued differentiation to Stage 4, they were harvested at the end of Stage 4. During harvest, cells were treated for 10-12 min with TrypLE at 37 °C. Released single cells were rinsed with DMEM/F12 and spun at 300 rcf for 5 min.

PP cells were further differentiated toward PDOs (S5-S7) by a slightly modified version of the published protocol ^78^. We added a step (Stage 5) by employed AggreWell™400 (STEMCELL, Cat#34415) plates to form controlled aggregates before embedding in the hydrogels. The plates were prepared prior to seeding following the manufacturer’s instructions. Single PP cells were seeded with a density of 200 cells/microwell (1.2x10^6^ cells/well in 6 well plate), then allow them to compact and well-form for 48 h. Cells at this stage are cultured under differentiated medium with cytokines and growth factors as described by Breunig *et al.,* without 5% Matrigel supplementation. Briefly, basal media BE3 containing MCDB131 (Thermo) with 2 mM L-Glutamine, 1.754 g/l Sodium bicarbonate, 0.44 g/l glucose (Sigma), 0.5% ITS-X (GIBCO), 44 mg/l L-Ascorbic acid and 2% fatty acid free BSA was dissolved and filtered through a 0.22 µm filter. PPs were formed as cell aggregates in basal media BE3 supplemented with 10 mM nicotinamide, 10 µM ZnSO4 (Sigma), 10 µM ROCK inhibitor, 50 ng/ml EGF, 50 ng/ml FGF10, 50 ng/ml KGF, and 50 nM MSC2530818 (Selleckchem) (named Stage 5-6 complete medium).

### 4.3 Aggregates were harvested from plates for further generation of PDOs in 3D hydrogel microenvironment

We sought to develop a matrix for the efficient generation of PDOs by recapitulating key physical and biochemical characteristics of the pancreatic microenvironment as an alternative to the established natural matrices Matrigel. To this aim, integrating both natural polymers and synthetic cross-linkers, dSIS-NB hydrogel was synthesized as described previously.

dSIS-NB hydrogels were crosslinked by 1.2 wt% dSIS-NB, 0.2 or 1.2 wt% PEG-4SH, and 6mM LAP. Aggregates collected from AggreWell™ microwells were encapsulated into either dSIS-NB hydrogel or undiluted Matrigel to generate PDOs. To create comparable conditions with equally sized PDOs, all precursor components were mixed prior to adding the aggregates to a density of approximately 3 × 10^3^ aggregates/mL. Aggregate-containing precursor solutions were mixed gently and pipetted 100 μL to create a round-shaped gel droplet into each 24-well glass bottom cell culture plate (NEST, Cat# 801006). The hydrogels were polymerized under 365 nm light (5 mW/cm^2^) for 2 min when encapsulated in dSIS-NB or incubated for 20 min at 37 °C when encapsulated in Matrigel.

After gelation, 1.5 mL of Stage 5-6 complete medium was added to each well. The medium was then refreshed on day 1 and day 3 post-encapsulation to induce pancreatic ductal precursors. For the induction of maturing PDOs, organoids were exposed to Stage 7 complete medium, which is made by supplementing basal medium BE3 with 10 mM nicotinamide, 10 µM ZnSO4, 50 ng/ml EGF, 50 ng/ml FGF10. The medium was added on day 5 post-encapsulation then changed every 2-3 days for 7-10 days. The differentiation process was monitored, and samples were collected at key stages.

### 4.4 Characterization of dSIS-NB hydrogel

dSIS-NB hydrogels were crosslinked by thiol-norbornene photo-click reaction. To prepare the hydrogel samples, 40 μL of the precursor solution (1.2 wt% dSIS-NB, 0.2 or 1.2 wt% of PEG4SH, and 6 mM of photoinitiator LAP) were dispensed in between two glass slides separated with 0.8 mm Teflon spacers and treated with a water-repellent coating. The assembly was placed under 365 nm UV light at 5 mW cm−2 for 2 minutes to yield dSIS-NB hydrogels with an approximate diameter of 8 mm and a thickness of 0.8 mm. Bulk hydrogel moduli were characterized by a modular rheometer (MCR 102, Anton Paar) operating in a strain-sweep mode with normal force (NF) control. The rheometrical testing was conducted at 25 °C, with a normal force of 0.25 N, shear strain from 1% to 5%, and a frequency of 1 Hz. The shear modulus (G’) in the linear viscoelastic region (LVE) was reported as the stiffness of the hydrogels.

The apparent morphology of dSIS-NB and Matrigel was observed by a scanning electron microscope (SEM) was used to show the microstructure of dSIS and dSIS-NB hydrogels. Bulk hydrogels were fixed in 0.1% fixing solution then washed in deionized water for 30 min three times to remove salt and residual chemicals before freezing at −80 °C and lyophilizing for 24 h under 20 Pa, −50 °C. These samples were placed in conductive adhesives and sprayed gold with an Emitech K575X sputter coater (Quorum Technologies Ltd., United Kingdom). Microstructural images were acquired using a JEOL JSM-7800F.

### 4.5 Morphological Analysis

The morphology of PDOs was assessed using brightfield microscopy and time-lapse imaging. The size, shape, and organization of the organoids were evaluated at different time points during the differentiation process. Images were analyzed using ImageJ software.

Phase-contrast images were acquired on a Nikon TE300 microscope with a 4× or 10× objective and a DS-FI2 camera. Confocal images were acquired with the BC34 (Oxford Instruments) confocal microscopes. Images were acquired with a 20× objective or a 10× objective.

Time-lapse imaging was achieved by maintaining the PDO culture plate in a VWR Water Jacketed CO2 Incubators (VWR International, LLC) under 5% CO2, 85% humidity, and 37°C. Bright field images were taken with a 4× or 10× objective in a CELLCYTE X™ live-cell imaging system (Cytena).

### 4.6 Gene expression analysis

At key stages, differentiated cells were collected for gene expression. Total mRNA was isolated by a NucleoSpin® RNA kit (MACHEREY-NAGEL) according to the manufacturer’s instructions. The mRNA concentrations were measured by A DS-11FX+ spectrophotometer (DeNovix, USA). OneLµg purified mRNA was used for each cDNA synthesis step. Real-time quantitative PCR (qRT-PCR) was performed using the TB Green Premix Ex Taq II kit (Takara, Cat# RR820L) with gene-specific primers listed in **Supplementary Table S3**. All gene expression studies were performed with three biological and three technical replicates for each experimental condition. To adjust for variations in the cDNA synthesis, data were normalized to GAPDH reference gene expression and presented as fold expression relative to control by 2^−ΔΔCt^ method.

### 4.7 Whole-mount immunostaining

PDOs were visualized by IF staining 12-day post-encapsulation. For whole-mount immunostaining, PDOs-laden hydrogels were fixed in 4% paraformaldehyde (PFA) at room temperature for 30 min. The hydrogel samples were then permeabilized in PBS with 0.3% Triton X-100 for 15 min at room temperature. Samples were then blocked for 1-2 h with 5% BSA in 0.3% Triton-X and stained with primary antibodies diluted in blocking buffer overnight at 4°C. Organoids were washed three times with PBS at 30-min intervals before secondary antibody staining for 2h at room temperature or overnight at 4°C. After a brief wash with PBS, nuclei were stained with 10 μg/mL DAPI (Sigma, Cat# D9542) in PBS for 10-15 min and then washed three times with PBS at 15-min intervals. The antibodies used are detailed in **Supplementary Table S4**. IF imaging was performed using a bench-top confocal microscope (BC43, Oxford Instrument). All images presented show representative results obtained from at least three independent experiments.

### 4.8 Flow cytometry

Cell monolayers at the end of stage 4 were dispersed as single cells using TrypLE™ for up to 15 min at 37 °C. Cell layer was gently triturated with a p1000 pipette. Once most cells were dispersed, double the volume of E8 medium was added to dilute TrypLE™, and the cell suspension was passed through a 40 µm strainer. The single cells were spun (3 min, 1000 rpm), resuspended in PBS, and cell count and viability were determined. Cells were stained for viability with LIVE/DEAD™ fixable near IR death cell staining kit (Invitrogen, Cat# L34976). After washing, the cells were fixed using Foxp3/ transcription factor buffer set (eBioscience, 00-5523-00) following the manufacture instructions. The fixed cells were stained with primary antibodies against PDX1 and NKX6.1, followed by the corresponding secondary antibodies listed in **Supplementary Table S4**. The stained cells were subjected for flow cytometry analysis using Attune NxT Acoustic flow cytometer (Thermo Fisher Scientific) and the data were analyzed using FlowJo™ v10 software (BD Life Sciences).

### 4.9 scRNA-seq library construction and sequencing

To isolate single cells from PDOs, samples from each gel condition were first dissociated in collagenase I. Cells from three gels from three different batches were pooled. The isolated PDOs were resuspended in TrypLE (Gibco) supplemented with 10nM Y27632 and incubated for 5 min in a water bath at 37°C. Cell dissociation was quenched by adding an equal volume of DMEM/F12 supplemented with 10% FBS. After centrifugation at 300 x g for 5 minutes, the cell pellets from each condition were processed according to manufacturer’s recommendations (Fluent BioSciences). Briefly, cells were added into Pre-templated Instant Partitions (PIPs) to capture mRNA during downstream process. The PIPs/cell emulsions capturing the mRNA were visually inspected upon manufacturer’s guideline to ensure their quality before proceeding with the next steps. Samples that met the standards of the PIPseq T2 v4.0PLUS Single Cell RNA Kit protocol were processed following the basic workflow of mRNA isolation, cDNA generation, cDNA amplification, library preparation, library pooling, and sequencing. The cDNA libraries were sequenced on a NovaSeq X Plus PE150 instrument at the Novogene (CA, US).

### 4.10 scRNA-seq processing

Raw sequencing reads were processed using PIPseeker™ (v3.3, Fluent BioSciences) with default parameters. Reads were aligned to the GRCh38 reference genome to generate gene–cell count matrices. Downstream analysis was performed using the Seurat (v5.3.0) R package.^79^ Quality Control and Doublet Removal Cells were retained for analysis based on the following quality criteria: (1) detection of between 3,000 and 50,000 unique molecular identifiers (UMIs); (2) expression of between 200 and 6,000 genes; and (3) mitochondrial gene content of <20%. To rigorously identify and remove artifacts, putative doublets were predicted and excluded using DoubletFinder (v2.0), assuming a doublet formation rate proportional to the cell loading density. Normalization, Batch Correction, and Clustering To normalize the data and mitigate technical variability, we applied SCTransform (v2), a regularized negative binomial regression method. This step modeled technical noise and regressed out confounding variables, including sequencing depth and percentage of mitochondrial reads. To ensure optimal handling of potential batch effects between samples (S7-Matrigel, S7-dSIS-NB, S6-dSIS-NB, and S5-Aggrewell), we performed a parallel comparative assessment using Harmony integration. However, the SCTransform workflow alone was found to sufficiently correct technical batch variation while preserving biological heterogeneity more faithfully than aggressive integration methods. Therefore, the SCTransform-normalized data were utilized for all subsequent dimensionality reduction steps. Principal Component Analysis (PCA) was conducted on the SCT-transformed residuals. The top 30 principal components (PCs) were selected to construct a Shared Nearest Neighbor (SNN) graph, followed by Louvain clustering at a resolution of 0.5 and Uniform Manifold Approximation and Projection (UMAP) for visualization. Differential Expression Analysis For cell type annotation and comparative analysis between conditions (e.g., S7-Matrigel vs. S7-dSIS-NB), differentially expressed genes (DEGs) were identified using the FindAllMarkers in Seurat with the Wilcoxon Rank Sum test. Genes were considered significantly differentially expressed if they met the following strict thresholds: (1) a natural log fold change (avg_log2FC) > 0.25; (2) expression detected in at least 10% of cells in either of the two groups comparison (min.pct = 0.1); and (3) a Bonferroni-adjusted p-value < 0.05.

### 4.11 Pseudotime and trajectory analysis

Differentiation trajectories were modeled by projecting cells of interest into a diffusion map space using the top 25 principal components (PCs) via the destiny R package (v3.1.1). The first two diffusion components were subsequently used as input for lineage inference using Slingshot (v2.16.0). The LGR5⁺ pancreatic progenitor cluster (Cluster 2) was explicitly defined as the root state for trajectory calculations. Principal curves representing the inferred lineage structures were projected into the diffusion space to visualize the developmental progression from progenitors to differentiated ductal cells.

### 4.12 Pseudobulk differential expression analysis

To perform robust differential expression analysis robust to single-cell noise, single-cell gene counts were aggregated to the sample level (pseudobulk) using Seurat (v5.3.0) and Matrix (v1.7-4). Genes with fewer than 10 accumulated counts across all samples were excluded from the analysis. Normalization and differential expression testing were conducted using the edgeR (v4.6.3) and limma (v3.64.3) packages following voom transformation. Differentially expressed genes (DEGs) were defined based on an absolute log₂ fold change ≥ 0.25 and a Benjamini–Hochberg adjusted *p*-value < 0.05.

### 4.13 Functional Enrichment Analysis (GO, KEGG, and FGSEA)

Gene Ontology (GO) enrichment analysis of identified DEGs was performed using clusterProfiler (v4.12.0) with the org.Hs.eg.db (v3.21.0) annotation database. Enriched terms across Biological Process (BP), Cellular Component (CC), and Molecular Function (MF) categories were identified using an adjusted *p*-value threshold of < 0.05. KEGG pathway enrichment was similarly performed using KEGGREST (v1.48.1). For rank-based enrichment, Fast Gene Set Enrichment Analysis (FGSEA) was conducted using the fgsea package (v1.34.2). Genes were ranked by a metric calculating the signed log₁₀(*p*-value) multiplied by the sign of the log₂ fold change derived from the differential analysis. Enrichment was assessed against the MSigDB Hallmark gene sets (v2023.1), with significance defined as an adjusted *p*-value < 0.05.

### 4.14 Gene Set Variation Analysis (GSVA)

To evaluate pathway activity changes across pseudotime or distinct cell states, GSVA was performed using the GSVA package (v2.2.0). “GO Biological Process” gene sets were used as input to compute pathway enrichment scores for individual cells (or pseudobulk samples) using a non-parametric kernel estimation. Differential GSVA scores between experimental groups were assessed using linear models in limma, and results were visualized using pheatmap (v1.0.13).

### 4.15 R package summary

The following R packages were used: Seurat (v5.3.0), SeuratObject (v5.2.0), harmony (v1.2.0), dplyr (v1.1.4), ggplot2 (v4.0.0), edgeR (v4.6.3), limma (v3.64.3), DoubletFinder (v2.0), GSVA (v2.2.0), fgsea (v1.34.2), clusterProfiler (v4.12.0), destiny (v3.1.1), slingshot (v2.16.0), TrajectoryUtils (v1.16.1), and pheatmap (v1.0.13). Gene annotation and pathway analyses utilized org.Hs.eg.db (v3.21.0), reactome.db (v1.92.0), KEGGREST (v1.48.1), and AnnotationDbi (v1.70.0). Supporting Bioconductor infrastructure included SingleCellExperiment (v1.30.1), SummarizedExperiment (v1.38.1), Biobase (v2.68.0), GenomicRanges (v1.60.0), GenomeInfoDb (v1.44.3), IRanges (v2.42.0), S4Vectors (v0.46.0), BiocGenerics (v0.54.0), MatrixGenerics (v1.20.0), and matrixStats (v1.5.0). Data manipulation and matrix operations used Matrix (v1.7-4), data.table (v1.17.8), generics (v0.1.4), and sp (v2.2-0).

### 4.16 Statistical analysis

Statistical analysis employed Prism 10 software (GraphPad Software, Boston, MA, USA). Results were expressed as mean ± SEM. Analysis between more than two sample groups was performed by a one-way ANOVA. Analysis between two samples was performed by a two-tailed unpaired Student’s t-test. A minimum statistical significance level of p L 0.05 was considered. Sample sizes are described in the figure legends.

### 4.17 Code and data availability

The scRNAseq analyses presented in the paper were performed with open-source algorithms as described in Methods. Further details will be made available by the authors on request. No custom code was generated during this study. The authors declare that all data supporting the findings of this study are available within the article, its Supplementary Information, attached files, and online deposited data (RNA-seq data -GEO number: GSE318769), or from the authors upon reasonable request.

## Author contributions

C.-C.L. and N.H.L. conceived the study and designed the research. N.H.L. performed the experiments and analyzed the data. V.T.D. and N.H.L synthesized dSIS-NB and conducted mechanical characterization of the dSIS-NB hydrogels. K.S., N.H.L., and X.B. performed scRNA-seq data analysis. N.H.L. wrote the original manuscript. C.-C.L. edited the manuscript. C.-C.L. also acquired funding and was responsible for project administration and resources. All authors reviewed and approved the final manuscript.

## Acknowledgement

This project was supported in part by the National Institutes of Health (R01DK127436).

