## Supplemental figures for "Photo-click Decellularized Matrix Hydrogels for Generating Pancreatic Ductal Organoids"

**Supplementary Figure Legends**

**Video S1. Disruption of the epithelial monolayer during early differentiation.** Real-time recording of cells during stage 2 differentiation showing breaks in the cell monolayer, illustrating dynamic changes in cell-cell and cell-ECM organization over time.

**Video S2.** **Morphology of pancreatic progenitor spheroids differentiated in Matrigel.** Time-lapse imaging (images were acquired every 2 h and displayed at 0.15 s per frame) of pancreatic progenitor spheroids cultured in Matrigel, demonstrating disorganized structures with limited lumen formation and extensive cellular spreading into the surrounding matrix.

**Video S3. Morphology of pancreatic progenitor spheroids differentiated in dSIS-NB hydrogels.** Time-lapse imaging (images were acquired every 2 h and displayed at 0.15 s per frame) of pancreatic progenitor spheroids cultured in dSIS-NB hydrogels (G′ ≈ 2.5 kPa), showing the formation of compact, cystic epithelial structures with well-defined boundaries.

**Video S4. Cytoskeletal organization of pancreatic progenitor spheroids in different matrices.** Confocal imaging of F-actin in pancreatic progenitor spheroids cultured in dSIS-NB hydrogels (left) and Matrigel (right), highlighting differences in epithelial organization and overall morphology between the two culture conditions.

**Video S5. Forskolin-induced swelling of pancreatic ductal organoids in different matrices.** Time-lapse imaging (Images were acquired every 30 min and displayed at 0.1 s per frame) of pancreatic ductal organoids cultured in dSIS-NB hydrogels (left) and Matrigel (right) following forskolin stimulation, illustrating matrix-dependent differences in organoid swelling responses.

**Supplementary Figures**

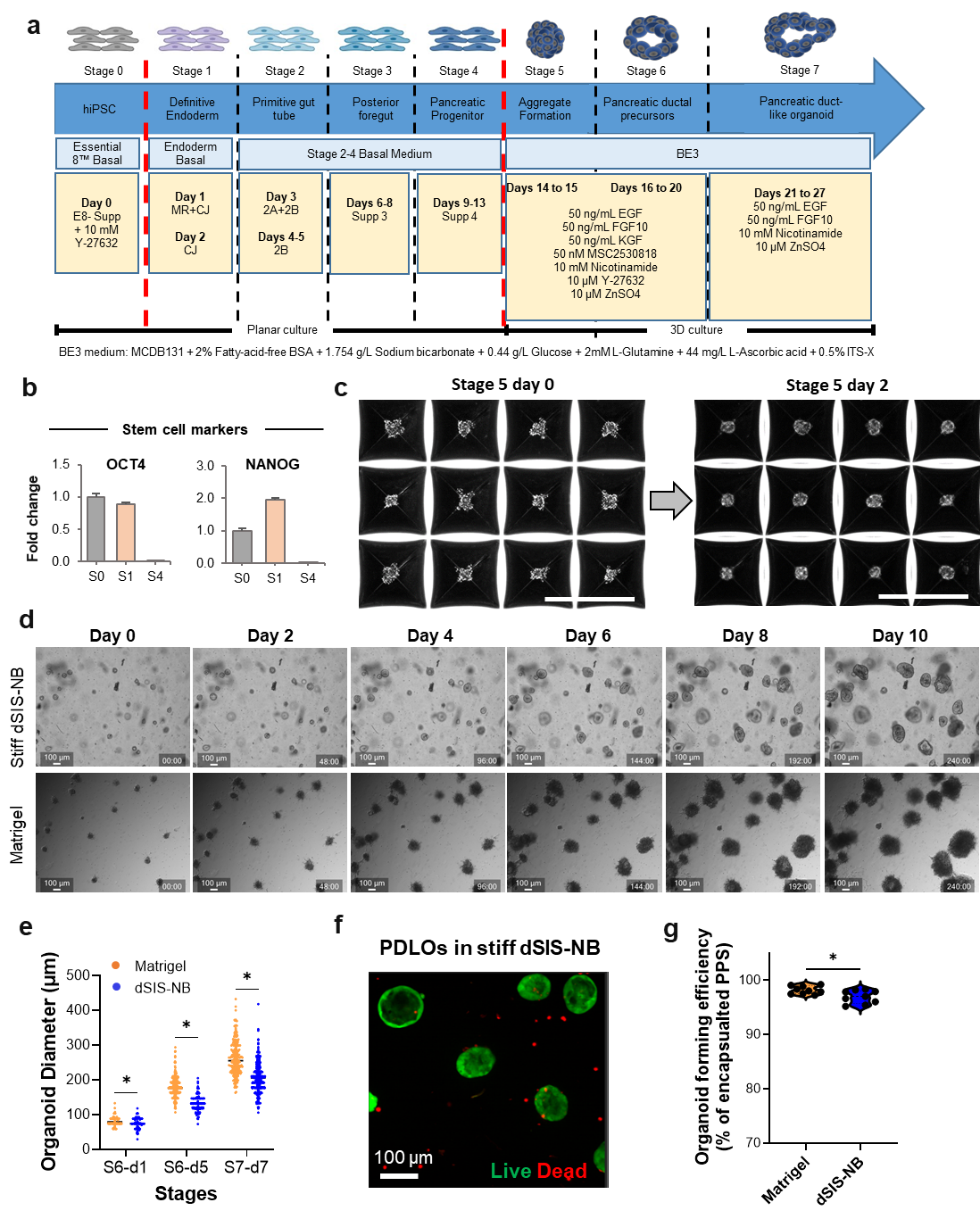

**Figure S1. Establishment and growth of PDOs in dSIS-NB hydrogels. a)** Schematic of the stepwise differentiation and organoid formation protocol from human iPSCs to pancreatic duct organoids (PDOs). Cells were guided through definitive endoderm, gut tube endoderm, pancreatic endoderm, and pancreatic progenitor stages (S0–S4), followed by 3D aggregation (stage 5) and maturation into pancreatic ductal precursors (stage 6) and PDOs (stage 7). Media compositions and timing are indicated. **b)** qRT-PCR analysis of stem cell-associated markers at the indicated stages. Expression of stem cell markers (OCT4 and NANOG) is enriched at early stages, whereas they are significantly downregulated at later stage (S4) (n = 3; mean ± SEM). **c)** Representative bright-field images showing aggregation of pancreatic progenitors at stage 5, imaged at day 0 and day 2 after aggregation, illustrating progressive compaction of cell clusters. Scale bars, 500 μm. **d)** Time-course bright-field images of organoid growth in stiff dSIS-NB hydrogels or Matrigel over 10 days following encapsulation. PDOs cultured in dSIS-NB progressively increase in size and display defined spherical morphologies, whereas Matrigel-cultured structures display disorganized morphologies with extensive cellular invasion into the matrix. Scale bars, 100 μm. **e)** Quantification of organoid diameter at indicated time points during stages 6 and 7 for PDOs cultured in Matrigel or dSIS-NB. Data points represent individual organoids (at least 255 organoids were measured for each condition, n = 3 biological replication); horizontal bars indicate mean values. **p* < 0.05. **f)** Live/dead staining of PDOs cultured in stiff dSIS-NB hydrogels, showing predominantly viable cells. Scale bar, 100 μm. **g)** Organoid-forming efficiency, expressed as the percentage of encapsulated pancreatic progenitor clusters that continuously outgrowth in Matrigel or dSIS-NB hydrogels. Data are shown as individual measurements with mean values indicated (n = 9); *P < 0.05.

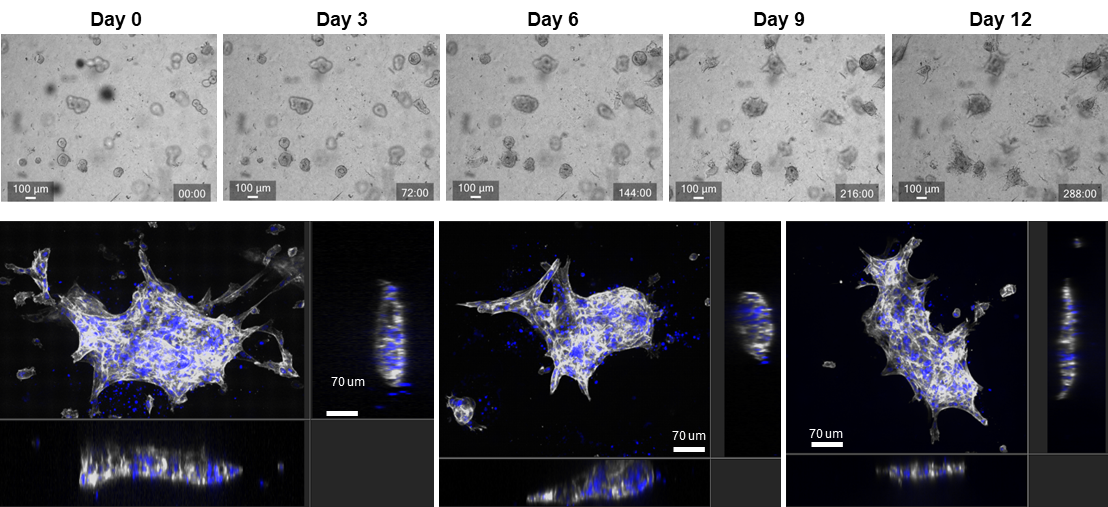

**Figure S2. PDO behavior in soft dSIS-NB hydrogels.** Representative time-course bright-field images of PDOs cultured in soft dSIS-NB hydrogels (0.2 wt% PEG4SH) over 12 days (0-288 h), showing progressive loss of spherical morphology. Scale bars: 100 μm.

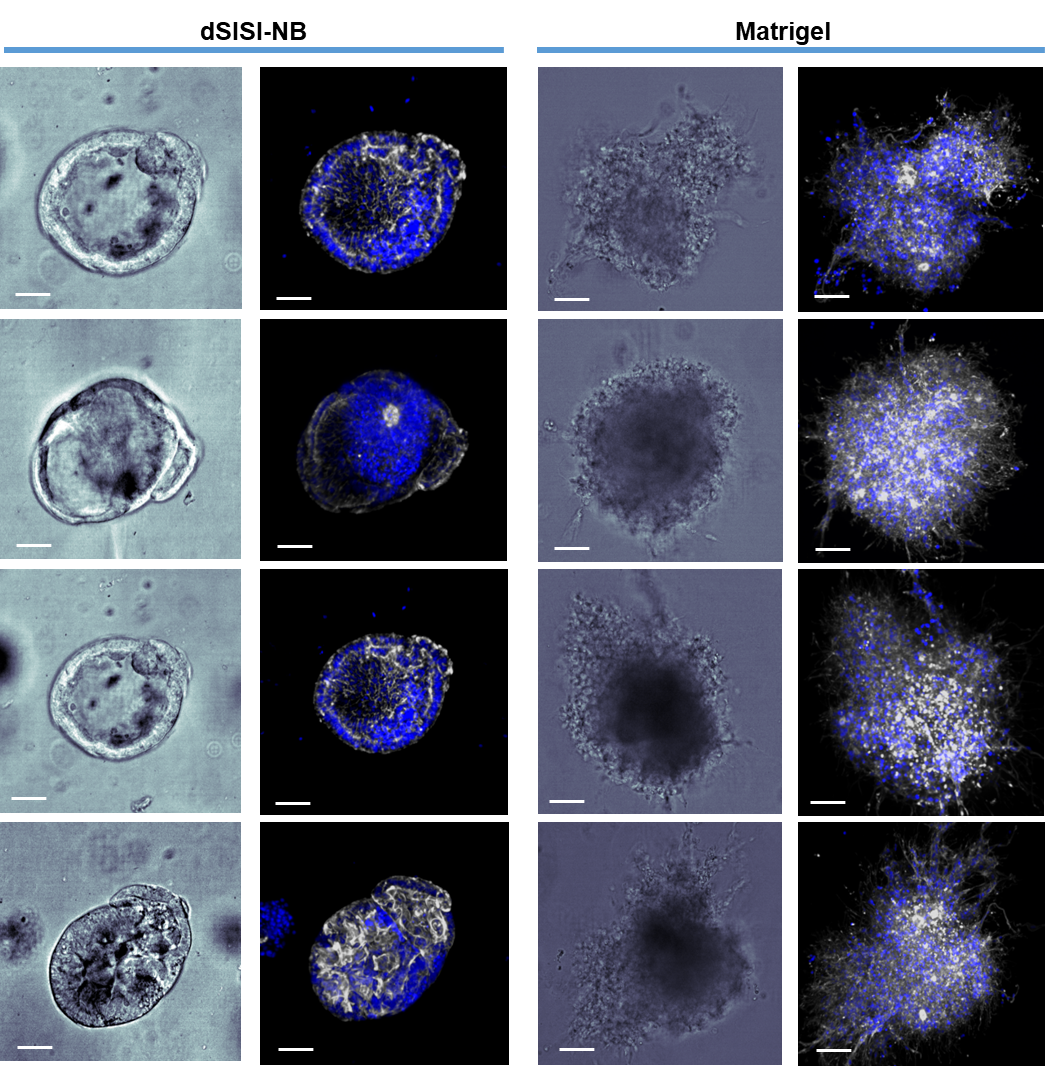

**Figure S3. Validation of dSIS-NB-mediated PDO morphogenesis using an independent iPSC line.** Representative bright-field and confocal images of PDOs derived from an independent WTC-mEGFP-CTNNB1-cl67 iPSC line (parent line hiPSC line: WTC/AICS-0) cultured in dSIS-NB hydrogels (left) or Matrigel (right). For each condition, bright-field images show overall organoid morphology, while corresponding confocal images display cytoskeletal organization and nuclear distribution (F-actin, white; nuclei, blue). PDOs cultured in dSIS-NB hydrogels consistently formed compact, rounded epithelial structures with defined boundaries across multiple examples, whereas those formed in Matrigel exhibited irregular, spread morphologies with extensive protrusive structures. Images are representative of multiple organoids analyzed across independent studies. Scale bars, 50 μm.

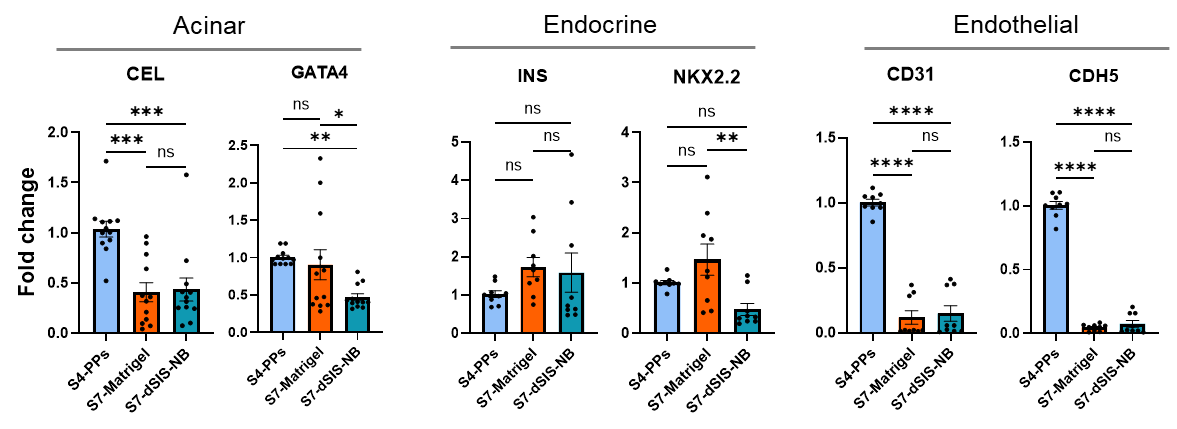

**Figure S4. Differentiation protocol limits off-target lineage gene expression.** Quantitative RT-PCR analysis of acinar (CEL, GATA4), endocrine (INS, NKX2.2), and endothelial (CD31, CDH5) marker genes in PDOs cultured in Matrigel or dSIS-NB hydrogels. Gene expression levels are shown as fold change relative to pancreatic progenitors (PPs). Across both culture conditions, expression of non-ductal lineage markers remained low, indicating minimal off-target differentiation. Data represent at least n = 3 independent differentiation batches and are presented as mean ± SEM. Statistical significance is indicated as ns, not significant; *P < 0.05, **P < 0.01, ***P < 0.001, ****P < 0.0001.

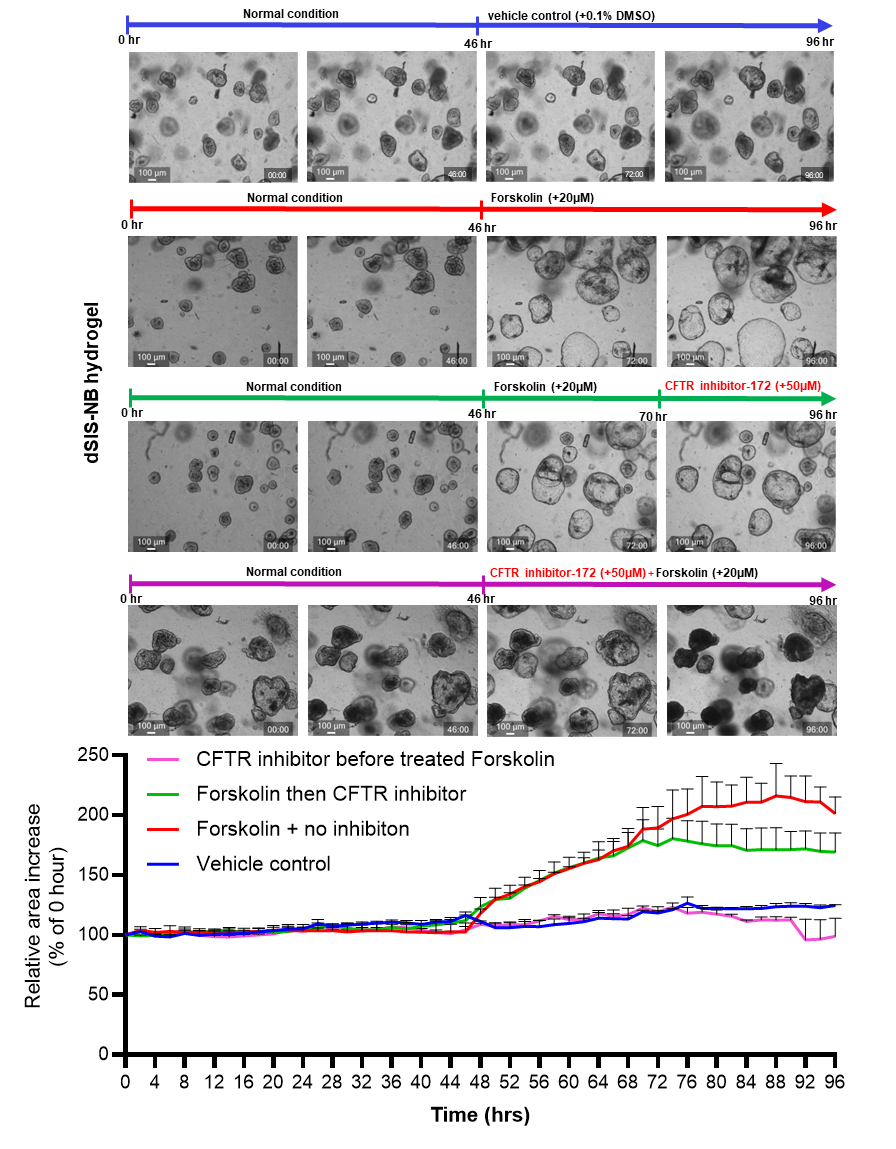

**Figure S5. CFTR inhibition attenuates forskolin-induced swelling in PDOs cultured in dSIS-NB hydrogel**. Bright-field images were used to quantify changes in PDO area following forskolin stimulation in the presence of a CFTR inhibitor. Area at each time point is expressed as percentage change relative to baseline (time 0). Measurements represent the average of two technical replicates per time point.

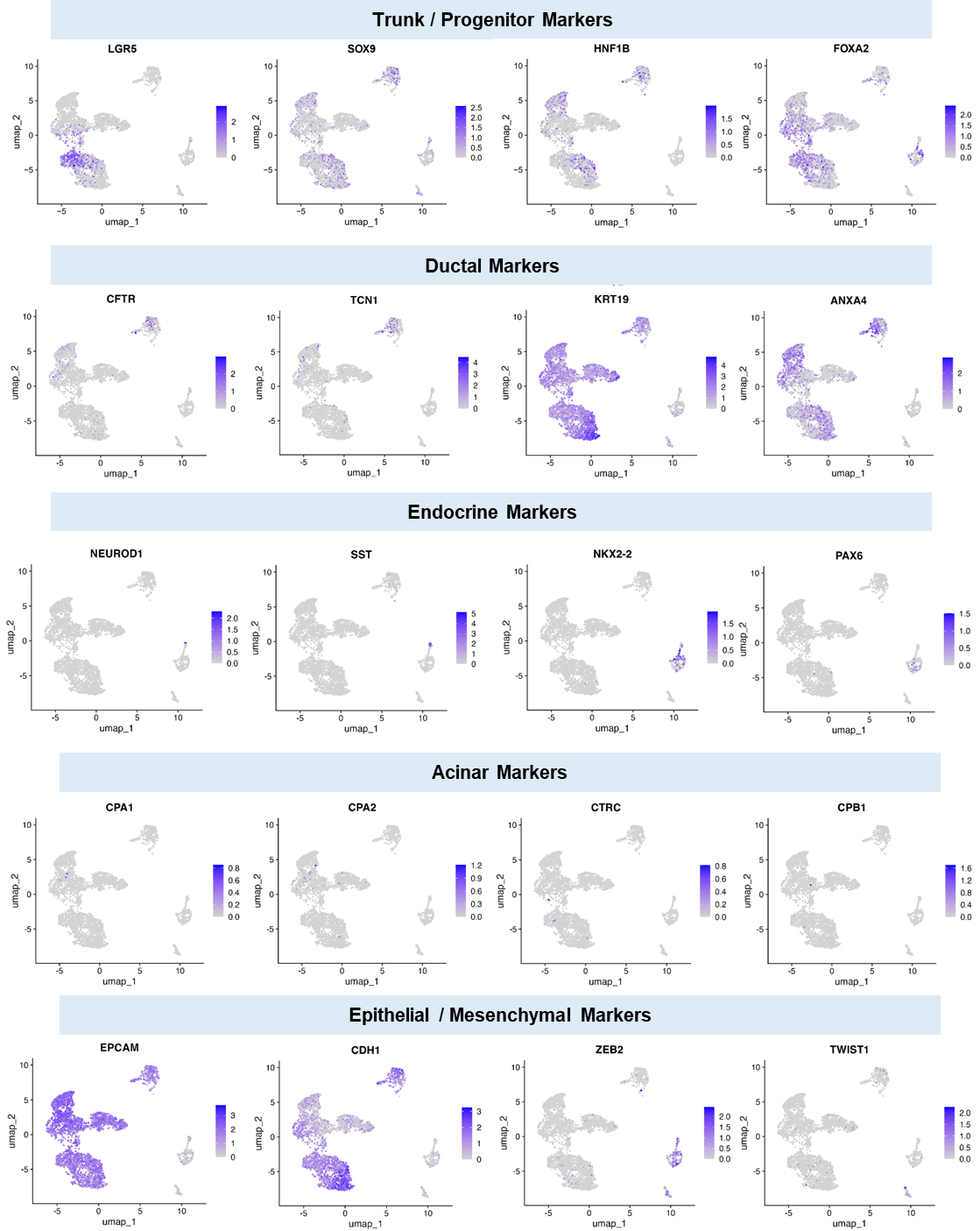

**Figure S6.** **UMAP feature plots of lineage and state marker gene expression.** Feature plots showing the expression patterns of representative genes grouped into five categories: Trunk/progenitor genes (LGR5, SOX9, HNF1B, FOXA2). Ductal genes (CFTR, TCN1, KRT19, ANXA4). Endocrine genes (NEUROD1, SST, NKX2-2, PAX6). Acinar genes (CPA1, CPA2, CTRC, CPB1). Epithelial and mesenchymal transition genes (EPCAM, CDH1, ZEB2, TWIST1). Color scales indicate log-normalized gene expression levels, with darker blue representing higher expression.

**Supplementary Tables**

**Table S1. Table S1. Differentially expressed genes between S7 PDLOs cultured in dSIS-NB and Matrigel (accessible in NCBI Gene Expression Omnibus site, GEO #GSE318769).**

**Table S2. Gene Ontology biological process enrichment analysis of pseudobulk RNA-seq data comparing S7 PDLOs cultured in dSIS-NB and Matrigel (accessible in NCBI Gene Expression Omnibus site, GEO #GSE318769).**

**Table S3. List of primers used for quantitative RT-PCR analysis**

| **Gene** | **Forward primer sequence (5’–3’)** | **Reverse primer sequence (5’–3’)** |
| --- | --- | --- |
| *GAPDH* | GCCTCCTGAAAAGAGAGTGGAAG | GCCTCCTGAAAAGAGAGTGGAAG |
| SOX9 | GCTCTGGAGACTTCTGAACGA | CCGTTCTTCACCGACTTCCT |
| NKX6-1 | CCTCATCAAGGATCCATTTTGTT | TGCTTCTTCCTCCACTTGGTC |
| PDX1 | ATGAACGGCGAGGAGCAGTA | TGGGTCCTTGTAAAGCTGCG |
| CFTR | AGAGGTCGCCTCTGGAAAAGG | GCGCTGTCTGTATCCTTTCCTCA |
| GATA4 | TCCCTCTTCCCTCCTCAAAT | TCAGCGTGTAAAGGCATCTG |
| NKX2.2 | GGTCCGGAGGAAGAGAACGA | AGACCGTGCAGGGAGTACTGA |
| INS | GAGGCTTCTTCTACACACCC | CCACAATGCCACGCTTCTGC |
| CEL | TGATGCTCACCATGGGGCGC | TGTACACGGCGCCCAGCTTC |
| KRT19 (CK19) | AACGGCGAGCTAGAGGTGA | GGATGGTCGTGTAGTAGTGGC |
| KRT7 (CK7) | TGTGGATGCTGCCTACATGAGC | AGCACCACAGATGTGTCGGAGA |
| CD31 | GCTGACCCTTCTGCTCTGTT | TGAGAGGTGGTGCTGACATC |
| CDH5 | AAGCGTGAGTCGCAAGAATG | TCTCCAGGTTTTCGCCAGTG |
| CDH1 | GTCCTGGGCAGACTGAATTT | GACCAAGAAATGGATCTGTGG |
| CDH2 | CCTCCAGAGTTTACTGCCATGAC | GTAGGATCTCCGCCACTGATTC |
| ZEB1 | AAGAATTCACAGTGGAGAGAAGCCA | CGTTTCTTGCAGTTTGGGCATT |
| VIMENTIN | CGAGGAGAGCAGGATTTCTC | GGTATCAACCAGAGGGAGTGA |
| EPCAM | GCCAGTGTACTTCAGTTGGTGC | CCCTTCAGGTTTTGCTCTTCTCC |
| SNAIL2 | GAAGATGCATATTCGGACCCACAC | TTGACCTGTCTGCAAATGCTCTGT |

**Table S4. List of antibodies used in this study.**

| **Antibodies** | **Dilution** | **Source/Isotype** | **Supplier (Catalog #)** |
| --- | --- | --- | --- |
| **Primary antibodies** |  |  |  |
| CK19 | 1-100 | Mouse | Santa Cruz Biotech (sc-53258) |
| CK7 | 1-100 | Goat | Santa Cruz Biotech (sc-17116) |
| SOX9 | 1-100 | Mouse | Santa Cruz Biotech (sc-166505) |
| CFTR | 1-400 | Rabbit | Cell Signaling (#78335) |
| SOX17 | 1-1000 | Rabbit | Cell Signaling (#81778) |
| PDX1 | 1-100 | Rabbit | Cell Signaling (#5679) |
| NKX6.1 | 1-100 | Mouse | DSHB (F55A12) |
| E-Cadherin | 1-1000 | Rabbit | Cell Signaling (#3195) |
| EpCam (CD326) | 1-100 | Mouse | Thermo Scientific (14-9326-82) |
| Vinculin | 1-100 | Mouse | Santa Cruz Biotech. (sc-25336) |
| Paxillin | 1-100 | Rabbit | Abcam (ab32084) |
| ZO-1 | 1-100 | Rabbit | Thermo Scientific (61-7300) |
| Ki67 | 1-200 | Rabbit | Cell Signaling (#9129) |
| Vimentin |  |  |  |
| **Secondary antibodies** |  |  |  |
| Anti-Rabbit Alexa Fluor 488 | 1-200 | Donkey | Invitrogen (A21206) |
| Anti-Mouse Alexa Fluor 488 | 1-200 | Goat | BioLegend (405319) |
| Anti-Rabbit Alexa Fluor 555 | 1-200 | Goat | Cell Signaling (#4413S) |
| Anti-Mouse Alexa Fluor 555 | 1-200 | Goat | Cell Signaling (#4409S) |
| Anti-Goat Alexa Fluor 555 | 1-200 | Mouse | Santa Cruz Biotech (sc-3916) |
